# Coinfections dampen the effects of temperature on host-parasite interactions

**DOI:** 10.64898/2025.12.10.693475

**Authors:** I. Will, C.A. Smith, T.E. Hector, K.C. King

## Abstract

Coinfections by multiple parasite species are widespread and modify disease outcomes. To what degree other forms of ecological complexity, such as environmental change, influence the impacts of coinfecting parasites on disease remains unclear. We exposed nematodes (*Caenorhabditis elegans*) to naturally coinfecting parasites (*Leucobacter celer* and *musarum*) at temperatures across the host’s thermal range. While we found that host mortality varied widely with temperature during single infections, coinfections buffered hosts against temperature-mediated variation via ongoing parasite competition. We extended our experiments with mathematical models indicating that changing virulence, rather than transmission, most plausibly underpins temperature-dependent host mortality. At moderate, but not extreme temperatures, we found coinfection continued to dampen mortality outcomes due to another source of host variation, genotypic susceptibility. Coinfection may provide a stabilizing effect on disease in a warming world – but this effect can falter when hosts are faced with multiple stressors.

## Introduction

Virtually all infected individuals are host to multiple parasite genotypes and/or species. Interactions between parasites can determine the level of virulence (i.e., host harm due to infection) as coinfections often modify disease outcomes in ways not predicted by one-host one-parasite paradigms (Alizon *et al*. 2013; Balmer & Tanner 2011; Graham 2008; Hoarau *et al*. 2020; Lello *et al*. 2018; Pedersen & Fenton 2007; Rigaud *et al*. 2010; Tollenaere *et al*. 2016). Exploitative parasite competition (e.g., for host resources) or facilitation (e.g., by modification of the immune system) can worsen disease for coinfected hosts (Alizon *et al*. 2013; Cressler *et al*. 2016; Ramesh & Hall 2025), increasing host mortality (Cornet & Sorci 2010; Louhi *et al*. 2015; O’Keeffe *et al*. 2024) and parasite burdens (Lass *et al*. 2013; Susi *et al*. 2015). Alternatively, parasites can diminish each other’s infectivity during competition thereby reducing the harmful effects of disease (Ford *et al*. 2016; Inglis *et al*. 2009; King *et al*. 2016). These “protective coinfections” can be conferred by apparent (e.g., via host immunity), exploitative (e.g., for public goods), or interference (e.g., by toxin production) forms of competition (Alizon *et al*. 2013; Ashby & King 2017; Cressler *et al*. 2016; Ramesh & Hall 2025). However, wild host organisms are exposed to variable and extreme environmental conditions likely to alter host resistance and tolerance, as well as parasite growth and transmission, that underpin disease outcomes.

Global climate change is predicted to promote novel or alter existing host-parasite and parasite-parasite interactions, driving emerging disease (Carlson *et al*. 2022; Morales-Castilla *et al*. 2021). The interaction of temperature and infection is often predicted to exacerbate disease during “thermal mismatches” where the host experiences more severe thermal stress than their parasites (Cohen *et al*. 2017, 2020; Sauer *et al*. 2020). Ectothermic animal hosts may be most vulnerable (Cohen *et al*. 2020), while smaller parasites may often gain the upper hand due to their broader thermal tolerances (Rohr *et al*. 2018), but certainly with exception (Gehman *et al*. 2018). The interaction of single species infections and temperature modifies both disease outcomes (Cohen *et al*. 2020; Hector *et al*. 2022; Ismail *et al*. 2024; Paull & Johnson 2014; Ware-Gilmore *et al*. 2021) and thermal performance (Bates *et al*. 2011; Hector *et al*. 2019, 2022; Hoang *et al*. 2019; Hudson *et al*. 2024; Martins *et al*. 2023; Wolinska & King 2009). Underpinning these processes, at least in part, the metabolic theory of ecology connects the basic kinetics of metabolism to organismal traits and interactions, such a parasitism (Brown *et al*. 2004; Kirk *et al*. 2018). This generalization can offer reasoned baseline predictions for host-parasite interactions, but may not well capture idiosyncratic responses or cases at thermal extremes (Kirk *et al*. 2018). Similarly building from fundamental principles, dynamic energy budget frameworks consider how resources are allocated (i.e., tradeoffs) within an individual and can help explain disease outcomes in light of the uptake and expenditure of energy (Douhard *et al*. 2025; Erazo *et al*. 2025; Hall *et al*. 2009). Temperature and disease could affect host uptake (e.g., food availability, foraging effort) and/or expenditure (e.g., immunity, thermal stress response, reproduction, and parasite consumption). However, these concepts have not been as widely and rigorously tested during coinfections, which are more relevant to wild disease ecology than single parasite infections (Telfer *et al*. 2010).

Environment, especially temperature, can modify coinfecting parasite prevalence (Fargues & Bon 2004; Friesen *et al*. 2021; Halle *et al*. 2024; Poterek *et al*. 2022), distribution (Oakgrove *et al*. 2014), growth (Budischak *et al*. 2015; Vaumourin & Laine 2018), and competition (Alcaide *et al*. 2021; Greenrod *et al*. 2024). Coinfection dynamics can reflect temperature impacts on individual parasite species occurrence (Halle *et al*. 2024) or more complex interactions stemming from asymmetric competitiveness mediated by thermal performance (Greenrod *et al*. 2024). The relationship of coinfecting parasites, host, and temperature has been shown to produce divergent virulence outcomes (Carpenter *et al*. 2024; Thomas *et al*. 2003) – although why, is not always known. Still unclear is if temperature dependent host-parasite effects modify coinfecting parasite interactions, thereby altering disease outcomes.

We tested whether coinfecting parasite interactions would persist across temperatures. Alternatively, temperatures that impact hosts could also impact coinfection interactions, altering disease outcomes. To test these hypotheses, we used naturally coinfecting parasites at temperatures representative of their source locality, covering a range of host thermal stress. We leveraged a nematode host (*Caenorhabditis elegans*) and two bacterial parasites (*Leucobacter celer* and *Leucobacter musarum*) co-isolated from a wild congener host, *Caenorhabditis tropicalis* (Bates *et al*. 2021; Clark & Hodgkin 2015). In *C. elegans*, *Leucobacter*-infected host mortality has been observed to vary across temperatures (*L. musarum*), moisture levels (*L. celer*), and host genotypes, and lead to tradeoffs between resisting either parasite (Bates *et al*. 2021; Hector *et al*. 2025; Hodgkin *et al*. 2013; Loer *et al*. 2015; O’Rourke *et al*. 2023; Parsons *et al*. 2014). *Leucobacter celer* attenuates or limits *L. musarum* infections during protective coinfection in the absence of extreme thermal stress (Bates *et al*. 2021; Hodgkin *et al*. 2013). We assessed host disease outcomes (mortality and fecundity loss) and parasite load across combinations of infections and temperatures spanning the host’s thermal optimum to upper reproductive limit. Thermal mismatches and metabolic theory would predict increasing parasite fitness moving from the lower host optimum temperature towards higher thermal stress for hosts, when these temperatures are well tolerated by the parasites (as in our experiments) (Cohen *et al*. 2020; Kirk *et al*. 2018). However, substantial changes in host energy budgets and physiology as a result of extreme thermal stress (i.e., castration) (Hall *et al*. 2009), may lead to deviations from those predictions. Based on these initial principles, we would predict that the fundamental relationship between temperature and disease to be similar for the congener parasite species tested. We further tested if parasite interactions were robust to direct (genetic) modification of host condition. Using mathematical models, we extended our experimental results to determine how transmission and virulence could explain temperature-dependent mortality, while remaining consistent with the effects of coinfection. Taken together, we tested how coinfection shapes disease when temperature radically alters host-parasite interactions.

## Methods

### Study system

We used the isogenic N2 strain *Caenorhabditis elegans* nematodes as hosts in all experimental assays. For mortality assays, we also used *srf-2* (*yj262*) mutant *C. elegans* (Politz *et al*. 1990). These *srf-2* mutants have a missense mutation in the *srf-2* gene (F59C6.8) producing cuticular glycosylation defects that impact locomotion and infection susceptibility (see below), but otherwise have near normal growth (O’Rourke *et al*. 2023). We maintained *C. elegans* on nematode growth medium plates (NGM) colonized by *Escherichia coli* (OP50) as a food source (Stiernagle 2006).

The two naturally co-isolated bacterial parasites we used were *Leucobacter celer* subsp. *astrifaciens* (CBX151) and *Leucobacter musarum* subsp. *musarum* (CBX152) (Clark & Hodgkin 2015). *Leucobacter* infections cause disease in both the wild host *Caenorhabditis tropicalis* (pers. comm. Serena Johnson and Luís Silva) and *C. elegans* (Hodgkin *et al*. 2013). *Leucobacter musarum* kills *C. elegans* in a temperature-dependent manner (Hector *et al*. 2025; Hodgkin *et al*. 2013). *Leucobacter celer* infection is essentially sub-lethal at those temperatures, but virulence has been measured by increased host generation times (Hodgkin *et al*. 2013) and gene expression indicating immunity-reproduction fitness tradeoffs (Will *et al*. 2025). Host tradeoffs between resisting one or the other parasite have been shown using *C. elegans* mutant lines (e.g., *srf-2*) (Bates *et al*. 2021; Hodgkin *et al*. 2013; Li *et al*. 2025; Loer *et al*. 2015; O’Rourke *et al*. 2023; Parsons *et al*. 2014).

To generate working cultures of *Leucobacter*, we inoculated 3 mL Lennox lysogeny broth (LB) from glycerol freezer stocks and incubated them at 25 °C for 24 h with shaking. We confirmed all cultures were within optical density (OD, 630 nm) 0.2 – 0.6 before proceeding with assay protocols. All OD measurements were taken in 96-well plates with 200 µl volumes with a Varioskan Lux plate reader (Thermo Scientific). For OP50, we instead inoculated 30 mL of LB, but otherwise followed the same methods.

### Parasite thermal optima

We estimated optimal growth temperatures for the parasites by combining *in silico* predictions and *in vitro* growth. We used Tome (v. 1.0.0) and its pre-trained machine learning model on proteome sequence data from both *Leucobacter* species to generate *in silico* predictions (Li *et al*. 2019). The predictions produced are a single point estimate of the optimal temperature, rather than performance across temperatures. For growth assays, we standardized cultures to OD 0.2 ± 0.02. Each biological replicate was a well-plate containing LB-only blanks (n = 12 wells) and *L. celer* and *L. musarum* inoculated samples (n = 24 technical replicate wells per species). Inoculated wells were filled with a 200 µL 1:10 dilution of initial OD 0.2 cultures in LB. Plates were then incubated with gentle shaking (60 rpm, “medium force”), with an OD reading every 15 min, for 24 h.

We conducted the growth curve assay at three temperatures – 20, 25, and 30 °C – with three biological replicate well-plates per temperature. The *Leucobacter* species were originally collected from Cape Verde, where soil temperatures range from approximately 15-30 °C and are trending upward (Hector *et al*. 2025; Lembrechts *et al*. 2022; World Bank Group & UEA Climatic Research Unit 2021). This temperature range includes thermal signposts for *C. elegans* reproduction: the optimum (20 °C), maximum (25 °C), and slightly beyond (30 °C) for *C. elegans* multi-generational fertility (Begasse *et al*. 2015), with some indication that the *Leucobacter* optimum is near 30 °C (Clark & Hodgkin 2015). We analyzed growth curve data using gcplyr (v. 1.10.0) in R to estimate maximum growth rates per well on median then average smoothed data (window = 7 data points for each) (Blazanin 2024).

### Infection mortality assays

We grew age-synchronized *C. elegans* using a bleaching protocol (Stevens *et al*. 2025). We plated approximately 1,500 larval stage 1 nematodes to 9 cm food plates and grew them at 20 °C for 48 h, until they were young adults, for subsequent experimentation. Mixed working cultures of bacteria were used to create infection assay plates of four kinds, all of which contained OP50 to feed the nematodes: OP50 only (control), *L. celer* (“LC” treatment), *L. musarum* (“LM” treatment), or both bacterial parasites (coinfection). All infection assay plates (6 cm NGM plates) received an inoculation of 80 µL of OP50 at OD 0.3. Of these plates receiving a parasite treatment (LM, LC and coinfection), we additionally added 20 µL per *Leucobacter* species at OD 0.2 to ensure that single-infection and coinfection treatments had the same initial inoculum per-species. All plates were incubated at 25 °C for 18 h to grow the bacteria to achieve consistent rates of environmental transmission to hosts. We then transferred approximately 100 of the young adult nematodes to each plate to begin the infection assay.

Our experimental design was fully factorial, combining infection treatments and environmental temperatures (Figure S1). Nematodes from each infection treatment or control treatment were placed at one of four temperature regimes for 30 h: 20 °C, 25 °C, 30 °C, or “heatwave”. Heatwave plates incubated at 25 °C for 12 h, then 35 °C for 4 h, before being returned to 25°C for 14 h. For Cape Verde (*Leucobacter* parasites source location), 35 °C soil temperatures have been recorded but are above average – suggesting that transient heatwave exposures are realistic (Hector *et al*. 2025). For the primary experiments with *C. elegans* N2, we conducted seven independent batches, each with two replicates (an assay plate with 100 nematodes) per treatment (n = 14 per treatment combination). To measure host mortality, we individually examined nematodes under a microscope; dead nematodes did not respond to gentle prodding by a wire. We conducted mortality counts aseptically for downstream host fecundity and parasite load assays.

To test if coinfecting parasite interactions continued to shape mortality outcomes in hosts with fundamentally altered susceptibilities to the parasites, we used mutant *srf-2 C. elegans* (see Study System, above). These mutants allowed us to manipulate host-parasite relationships without directly altering parasite-parasite relationships, facilitating a comparison between single-infection and coinfection processes. For *srf-2 C. elegans*, we conducted four batches of mortality assays, one with four replicates and the rest with two replicates per treatment (n = 10 per treatment combination) conducted in the same manner as N2 mortality assays.

### Parasite load and host fecundity assays

Using a subset of the *C. elegans* N2 infection assay plates, we quantified parasite loads (colony forming units, CFUs) (5 of 7 infection batches, n = 10 per treatment) and host fecundity (3 of 7 infection batches, n = 6 per treatment, a subset of the batches used for parasite load) (Figure S1). After scoring mortality, we used a wire to pick 13 live nematodes per plate and transferred them to tubes with M9 buffer (Stiernagle 2006) supplemented with 0.1% Triton-X-100 (TX). We washed the nematodes three times in M9-TX and suspended them in a final volume of 200 µL of M9. As *Leucobacter* adheres to the host surface (Hodgkin *et al*. 2013), we did not surface-sterilize nematodes (Figure S2). Of the 13 washed nematodes, we allotted 10 individuals to be used for CFU assays (see below) and three for fecundity assays (Supporting document S1).

We estimated *Leucobacter* loads by counting bacterial CFUs from infected hosts. We selected only live individuals for this assay. We aimed to capture parasite loads during infection of live hosts instead of consumption of dead individuals. Although these survivors may have avoided or tolerated infection to a greater degree than their dead counterparts within a replicate, they permitted more consistent comparisons across treatments. Using 10 washed nematodes picked from infection plates (see above), we added zircon beads (0.5 mm) to the nematode suspension and homogenized samples (Qiagen TissueLyser II, 5 min, 30/s frequency). We only did this for samples with *Leucobacter* infections. We serially diluted all samples to 1:1,000 in M9. We spotted three 10 µL technical replicates of each diluted sample onto LB agar plates and incubated them at 25 °C for 48 h. Colony size and color between *L. celer*, *L. musarum*, and OP50 are distinct and can be distinguished by eye. We recorded the number of parasite CFUs per pool and calculated the average per nematode. We verified that environmental competition (co-culture on assay plates) was likely only a minor influence on host bacterial load by measuring *Leucobacter* CFUs from infection assay plate lawns (Figure S3).

### Statistical models

We used generalized linear mixed models (GLMMs) to identify how infection and temperature may drive host mortality, host fecundity, and bacterial load. We analyzed host mortality with logistic regressions (binomial live and dead counts) using a Bayesian framework to better converge a model with cases of data separation (no death in 20 °C control samples). We evaluated effects by considering their posterior distribution 95% credible intervals (95CI). We set informed priors for Bayesian mortality models (see below) corresponding to a mortality probability distribution that reflects the range of mortality outcomes previously reported for *Leucobacter* infection of *C. elegans* (Bates *et al*. 2021; Hector *et al*. 2025). For modeling continuous responses (host fecundity and bacterial CFUs) that did not have data separation, we used frequentist negative binomial GLMMs as they were computationally expedient. We used a p ≤ 0.05 threshold for significance in frequentist tests. For all models, we considered temperature (levels: 20 °C, 25 °C, 30 °C, and heatwave) and infection (levels: control, LC, LM, and coinfection) as categorical fixed effects. For all models, we included experimental batch as a random intercept effect. We reported a Bayes R^2^ for Bayesian models and a Nakagawa’s pseudo-R^2^ for frequentist models. We conducted all statistics with R (v. 4.4.0) in RStudio (v. 2023.12.1) (R Core Team 2022; RStudio Team 2020). We used R packages brms (v. 2.21.0) for Bayesian models, glmmTMB (v. 1.1.9) for frequentist models, and performance (v. 0.11.0) to assess models (Brooks *et al*. 2017; Bürkner 2021; Lüdecke *et al*. 2021). For brms models, we set high run parameters with 10,000 iterations (2,000 warmup), 12 chains, 0.9999 adapt delta, and a max tree depth of 15. Our priors were “normal(-2, 3)” on the logit scale for the intercept and all treatment categories with the default “student_t(3, 0, 2.5)” for random batch effects. The negative binomial data family we used with glmmTMB was “nbinom2”. To extract predictions from model fits on the response scale, we used brms function conditional_effects including random effects (re_formula=NULL) for the mortality models and package ggeffects (v. 2.3.2) (Lüdecke 2018) function predict_response for the CFU models. We also used R package ggplot2 (v. 3.5.1) and Inkscape software (v. 1.3.2) to produce and finalize figures (Wickham 2016).

### Mathematical models

We used mathematical modeling to understand how infection virulence and parasite transmission could drive divergent host disease outcomes across conditions (temperature, in our system). We constructed the models to allow changes in both transmission and virulence to explain population-level mortality. For the mathematical models, we defined transmission to describe the transition from a susceptible host to infected host, rather than a rate of parasite accumulation or load. We defined virulence as mortality caused by infections. Rather than directly modeling temperature, our models described general-case, non-linear shifts in mortality, exemplified by those we observed from different temperature by parasite species interactions. We first described how virulence and transmission can shift mortality due to one parasite species in the environment (Figure 1A, Supporting document S2). We then constrained this virulence-transmission “mortality landscape” by the effect of coinfection. Coinfections encompass diverse interactions; here, we represented the effect of coinfection as intermediate mortality (i.e., protection) to align with our empirical findings (see Results). We considered an exclusion (transmission-reducing) mechanism and an attenuation (virulence-reducing) mechanism of protection (Figure 1B). In either case, we required model simulations to produce intermediate mortality during coinfection, thus constraining valid virulence-transmission values initially established by single infections.

**Figure 1.**
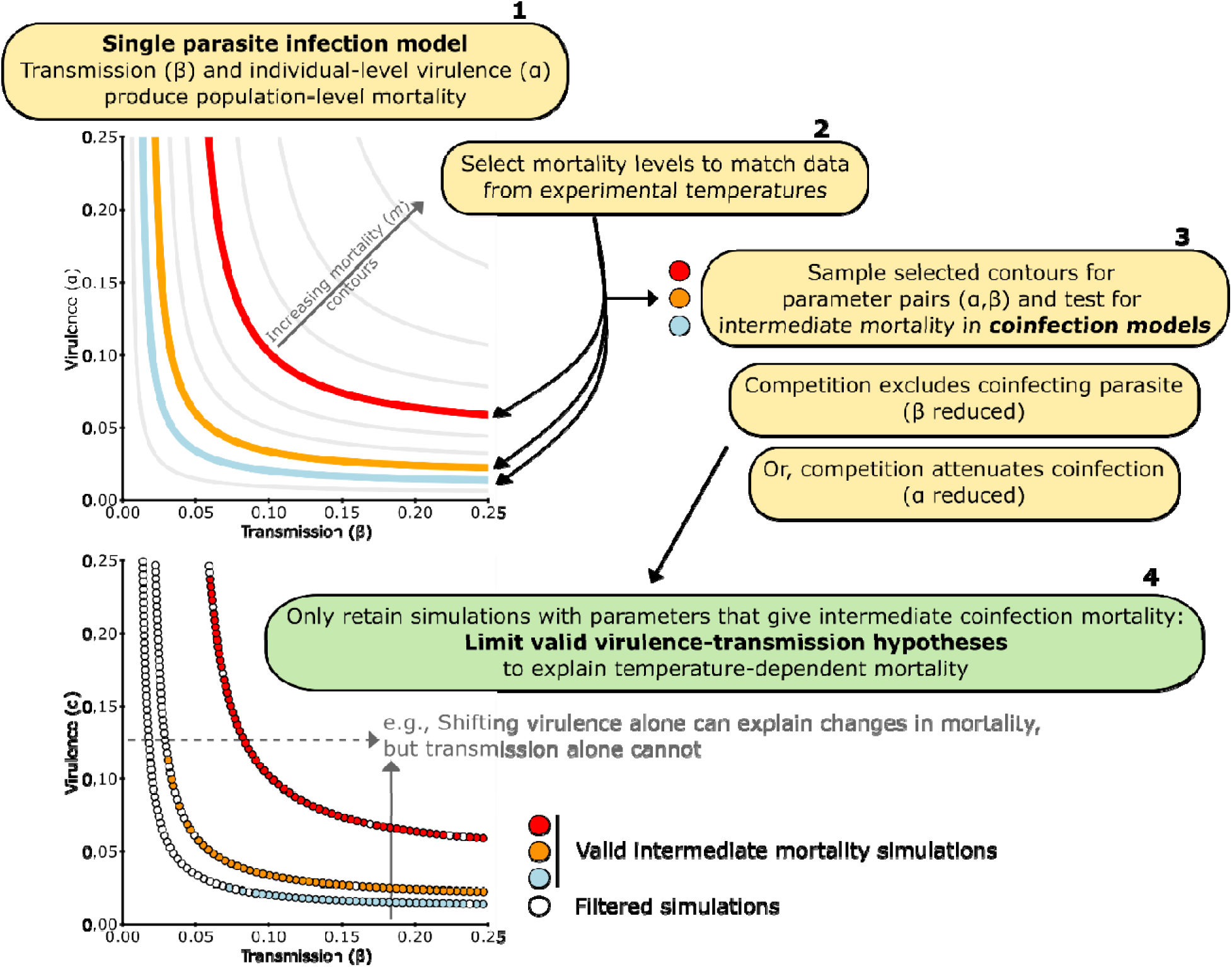
Coinfection was used to constrain single-infection simulations of host population mortality to understand possible temperature-virulence-transmission relationships. The initial mortality contours (each contour represents some mortality level, *m*) for a single infection created a theoretical mortality landscape that established the fundamental ways that transmission (β) and virulence (α) can drive population-level mortality (**1**). Contours farther from the axes indicate higher levels of mortality. We extracted virulence-transmission parameter pairs along selected contours, chosen to reflect patterns in our experiments (**2**). We then tested those parameter pairs in coinfection models of either exclusion (β reduced by ) or attenuation (α reduced by ) (**3**). We required that valid simulations produced intermediate mortality during coinfection (an effect found in our experiments) (**4**). The remaining truncated virulence-transmission contours constrained viable paths to move between mortality levels. Additional details can be found in Supporting document S2.

Here, we describe the key assumptions and governing equations of our mathematical models (Supporting document S2). We modelled a host population that can be infected by two parasites, indexed as 1 or 2, either singly or in coinfection. The parasites were environmentally transmitted (as *Leucobacter* are in our experiments), and so infection occurs with a flat rate *α_j_* (transmission) for parasite *j*= 1, 2, and parasites cause mortality at rate *β_j_* (virulence) in infected hosts. We assumed the timeframe of the simulation is shorter than host generation time (i.e., no host births). Non-disease induced mortality was included as rate d. Finally, we included parasite competition that produces intermediate mortality by either exclusion (reducing *β*) or attenuation (reducing *α*). The model did not distinguish if virulence attenuation might be driven by either resistance or tolerance. Exclusion was encapsulated by functions *E*_l_(*y_e_*_l_) (resp. *E*_2_(*y_e_*_2_)) for varying exclusion by parasite 1 (resp. 2) against parasite 2 (resp. 1) with strength *y_e_*_l_ (or *y_e_*_2_) (see equation below). We define the strength of exclusion conferred to a host with parasite 1 against parasite 2 as *y_e_*_l_ E [0, 1], where 0 is no exclusion and 1 is complete exclusion. Alternatively, we considered attenuation, given as functions *A*_l_(*y_a_*_l_) (resp. *A*_2_(*y_a_*_2_)), following the same naming convention. The ordinary differential equations corresponding to these assumptions are as follows, where we define *N*_O_ to be the initial population size, S(t) to be the density of the susceptible population, *l*_l_(*t*) and *l*_2_(*t*) to be the densities of the singly-infected populations, and C(t) to be the coinfected population density, after time some time *t*:

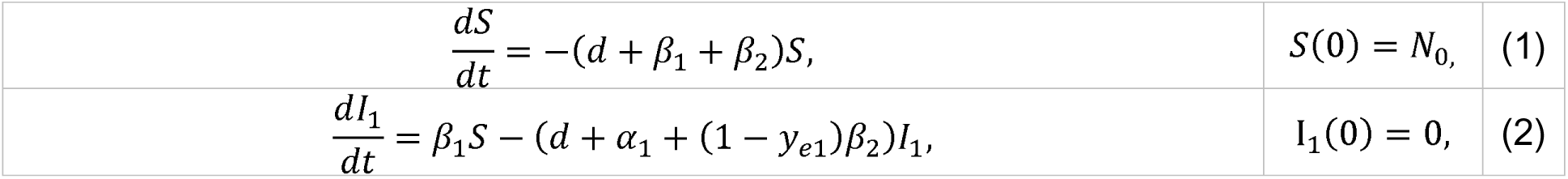
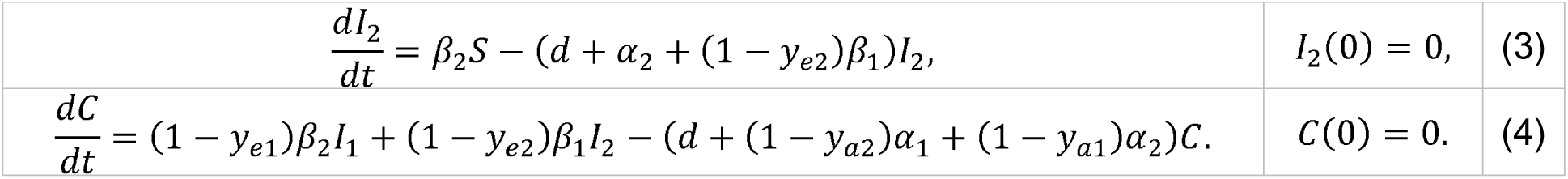

Finally, mortality, m = M(T), is defined as the total proportion of the initial proportion that has died at the final time, T (Supporting document S2). In our models, T was fixed arbitrarily to the same time in all simulations. When testing the effect of exclusion or attenuation, the effect of the other was set to zero for simplicity. The model and equations in their entirety can be found in the Supporting document S2.

We then used these models for simulations (Supporting document S2). For each temperature (i.e., mortality level), we sampled 1,000 parameter combinations for parasites 1 and 2 (*β*_l_, *α*_l_, *β*_2,_ *α*_2_). Per parameter combination, we then sampled 20 protection values (*y_e_*_l_, *y_e_*_2_) or (*y_a_*_l_, *y_a_*_2_) uniformly at random from the unit square, and calculated coinfection mortality (m) (Supporting document S2). From these coinfection mortalities, we filtered simulation results to satisfy the requirement of coinfection producing intermediate mortality between single infections. By plotting the accepted simulations, we ascertained how virulence and transmission could change across conditions (temperature) to produce mortality shifts exemplified by the patterns we observed experimentally.

## Results

### Parasites had a higher thermal tolerance than their host

Genomic predictions and *in vitro* growth curve assays indicated both parasites well tolerate 20-30 °C temperatures. Our computational analysis of the *Leucobacter* proteomes returned predicted thermal optima of 29.2 °C and 28.9 °C for *L. celer* and *L. musarum*, respectively. Corroborating those findings empirically, we measured the highest maximum growth rate for both species at 30 °C (2.3 and 2.0 h median doubling times for *L. celer* and *L. musarum*, respectively) as opposed to cultures grown at 20 or 25 °C (2.8-2.9 h doubling times) (Figure S4).

### Parasite fitness was reduced by coinfection across varying effects of temperature

To test whether temperature and coinfection interactions modified parasite fitness, we modeled parasite fitness (CFU loads) as “Load ∼ Temperature*Infection + (1|Batch).” We created two such GLMMs, one for *L. celer* and one for *L. musarum* loads, where the infection term could be either single or coinfection. For *L. celer* (Nakagawa’s pseudo-R^2^ = 0.601, conditional and 0.441, marginal), we observed effects of both temperature and coinfection, but not their interaction (Figure 2A, Figure S5, Table S1). For *L. musarum* (Nakagawa’s pseudo-R^2^ = 0.620, conditional and 0.479, marginal), we observed effects of both temperature and coinfection, and a specific interactive effect between coinfection and heatwave (Figure 2B, Figure S5, Table S2).

**Figure 2.**
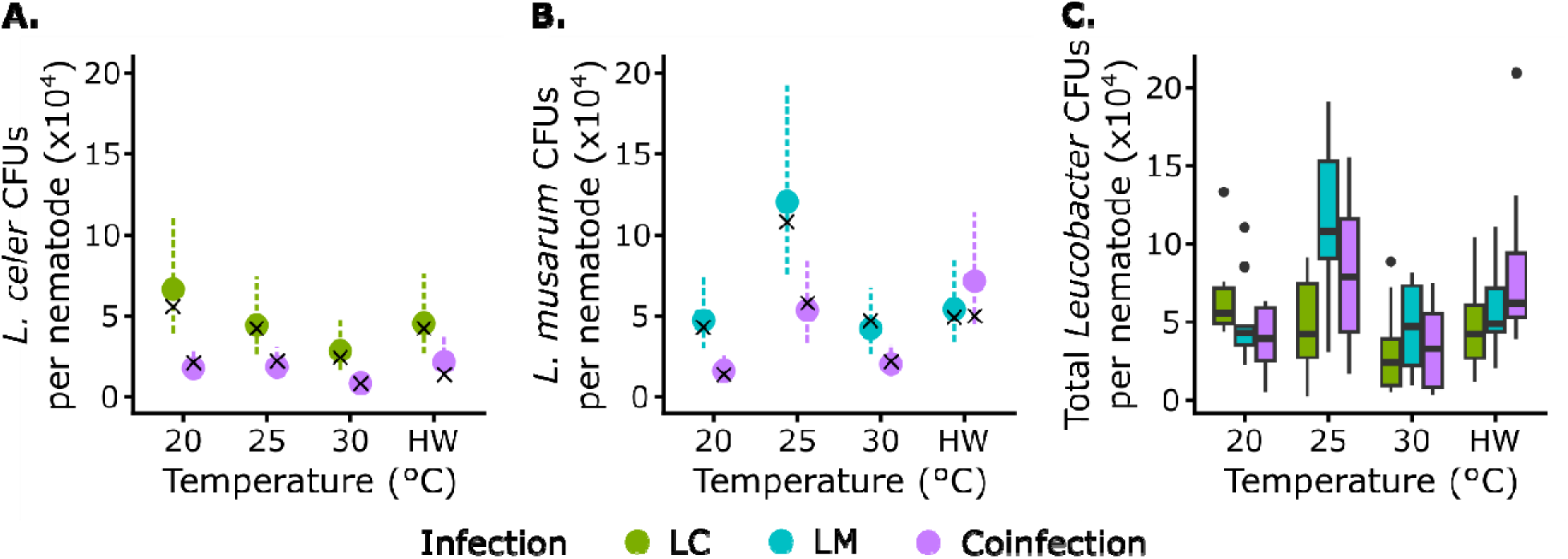
Parasite fitness (CFU load) was generally reduced by coinfection, with secondary effects of temperature. (**A.**) *Leucobacter celer* load was reduced by high temperature (30 °C) and by coinfecting *L. musarum* across temperatures. (**B.**) *Leucobacter musarum* load peaked at 25 °C and was typically suppressed by coinfection. However, during heatwave, *L. musarum* loads were not reduced by coinfection. Load reductions during coinfection were larger than observed during in vitro co-culture on assay plates alone (Figure S3). Per nematode values were calculated from 10-nematode replicate pools. Points are predicted values from the GLMM with frequentist 95% confidence intervals as dashed lines (rather than Bayesian credible intervals reported for mortality models). Median values of the raw data (Figure S5) are given by crosses. (**C.**) The boxplots show observed total parasite loads (i.e., coinfection treatments sum *L. celer* and *L. musarum* CFUs). Although coinfection reduced loads of both parasite species individually (A. and B.), the total *Leucobacter* load on coinfected hosts was not clearly below the lowest single-infection load at any given temperature (C.).

*Leucobacter celer* growth in hosts was suppressed at high temperature and by coinfection with *L. musarum*. Corresponding to peak LC treatment mortality (Figure 3), sustained high temperature at 30 °C reduced *L. celer* loads during single infection (effect = −0.85, z = −2.89, p = 0.004) (Figure 2A, Table S1). Coinfection further lowered *L. celer* loads at all temperatures, with a 61% reduction at 20 °C (effect = - 1.32, z = −4.48, p < 0.001) (Figure 2A, Table S1). We did not detect a clear interaction of coinfection and any temperature, indicating that the strength of parasite competition on *L. celer* was not sensitive to warming (Figure S8, Table S1).

**Figure 3.**
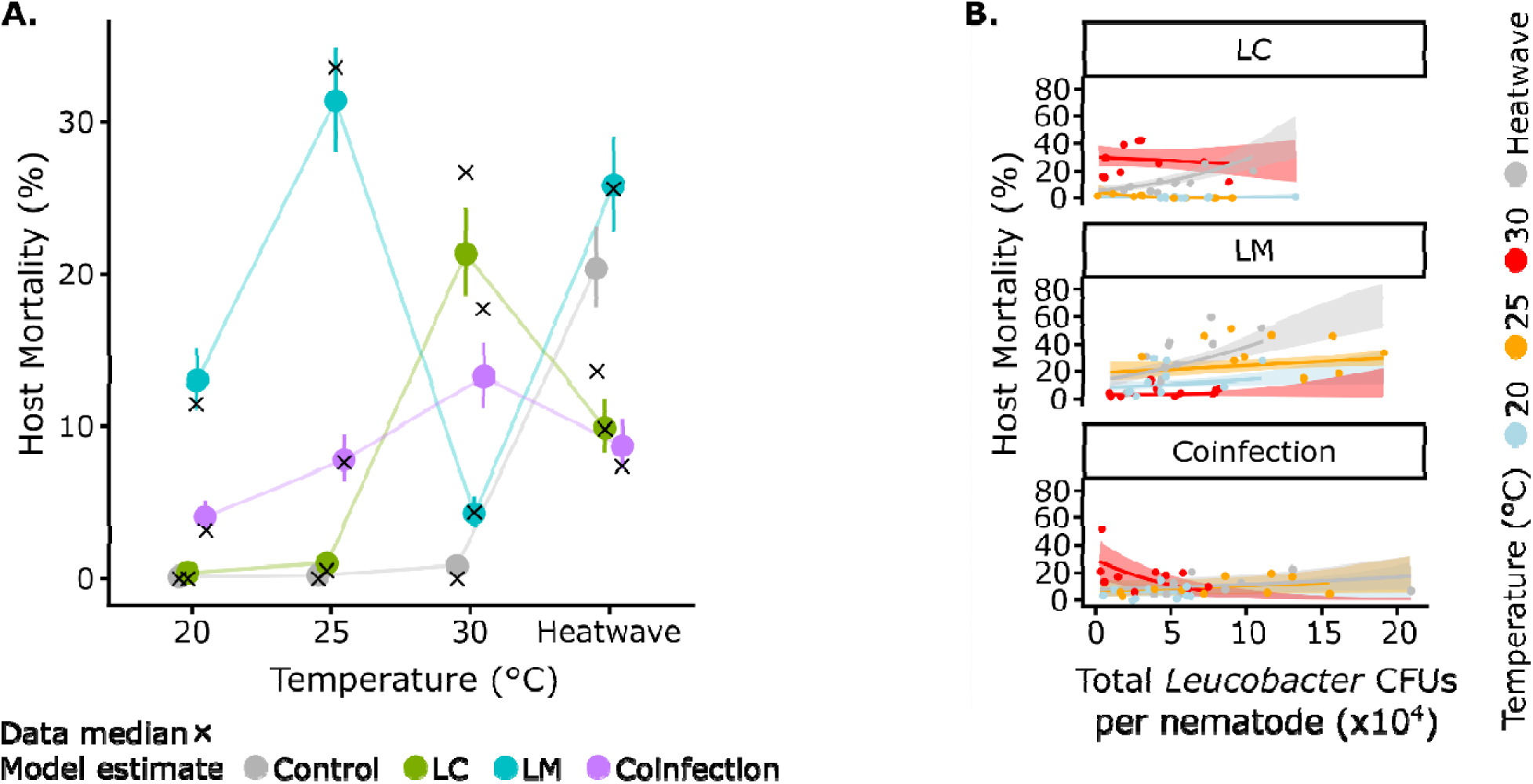
Coinfecting parasite interactions modified temperature-dependent host mortality. (**A.**) Model estimates (points) and 95CI (bars) show LM (*L. musarum* alone) and LC (*L. celer* alone) treatments with extreme warming (30 °C) reversed in relative severity. However, at all temperatures, coinfection protected hosts from peak mortality of the most harmful single infection. Median values of the raw data (Figure S6) are given by crosses, typically laying within the 95CI from model predictions. Points are connected by lines to show trends. (**B.**) Parasite loads and host mortality did not have consistent, strong trends correlating *Leucobacter* CFUs and host death. Only LM treatments showed positive relationships between load and mortality, although these effects were not very strong (Supporting document S3). LC infections during heatwave did not have sufficient statistical support to link load and mortality, despite an apparent positive slope (Supporting document S3). Plotted points are the observed data, model estimates are given by the solid line with 95CI shaded ribbons. The estimate solid lines are truncated at the maximum observed CFU value per treatment.

Within-host *L. musarum* loads were enhanced with moderate warming and typically reduced by coinfection. LM treated hosts had increased parasite loads at 25 °C (effect = 0.93, z = 3.54, p < 0.001) (Figure 2B, Table S2), where we observed peak LM host mortality (Figure 3). *Leucobacter musarum* abundance was reduced during coinfection at most temperatures, with a 67% decrease at 20 °C (effect = - 1.07, z = −4.10, p < 0.001) (Figure 2B, Table S2). However, during heatwave, *L. musarum* loads were not reduced by coinfecting *L. celer* (heatwave-coinfection interactive effect = 1.35, z = 3.63, p < 0.001) (Figure 2B, Figure S8, Table S2). This change in underlying competition dynamics led to coinfection maintaining total parasite loads at or above the highest single-infection during heatwave (Figure 2C). At all other temperatures, coinfection had loads near or slightly above the lowest single-infection loads (Figure 2C).

### Coinfection dampened the impacts of temperature on disease outcomes for hosts

To test to what extent temperature and infection could predict host mortality, we constructed a Bayesian logistic regression (Table S3). This GLMM can be summarized as “Mortality ∼ Temperature*Infection + (1|Batch)” with a conditional Bayes R^2^ of 0.75 and a marginal of 0.54. We found strong effects due to temperature, infection, and their interaction on mortality (Figure 3A, Figure S6, Table S3). We additionally constructed multiple GLMMs of host mortality using parasite load and temperature predictors (Figure 3B), detailed in Supporting document S3. Implicit in these models with load was the assumption that parasite load may drive host outcomes, rather than moribund hosts becoming differentially colonized.

Host mortality varied in a parasite species and temperature dependent manner. We observed the clearest differences between moderate temperatures (20-25 °C) and severe thermal stress (30 °C). At both 20 and 25°C, *L. celer* alone promoted negligible host mortality (0 and 0.5% LC median mortality, respectively) while *L. musarum* only infections worsened with mild warming (11 and 34% LM median mortality, respectively) (Figure 3A, Table S3, Supporting data S1) (Bates *et al*. 2021; Hector *et al*. 2025; Hodgkin *et al*. 2013). When temperature increased to 30 °C, we found opposing temperature-by-infection interactive effects between LC and LM treatments (Figure 3A, Figure 4, Table S3). LC treatment markedly increased host mortality relative to controls at 30 °C (27% median mortality) (Figure 3A, Figure 4, Table S3). Conversely, the LM treatment mortality was much reduced (4% median mortality) (Figure 4, Table S3) (Hector *et al*. 2025).

**Figure 4.**
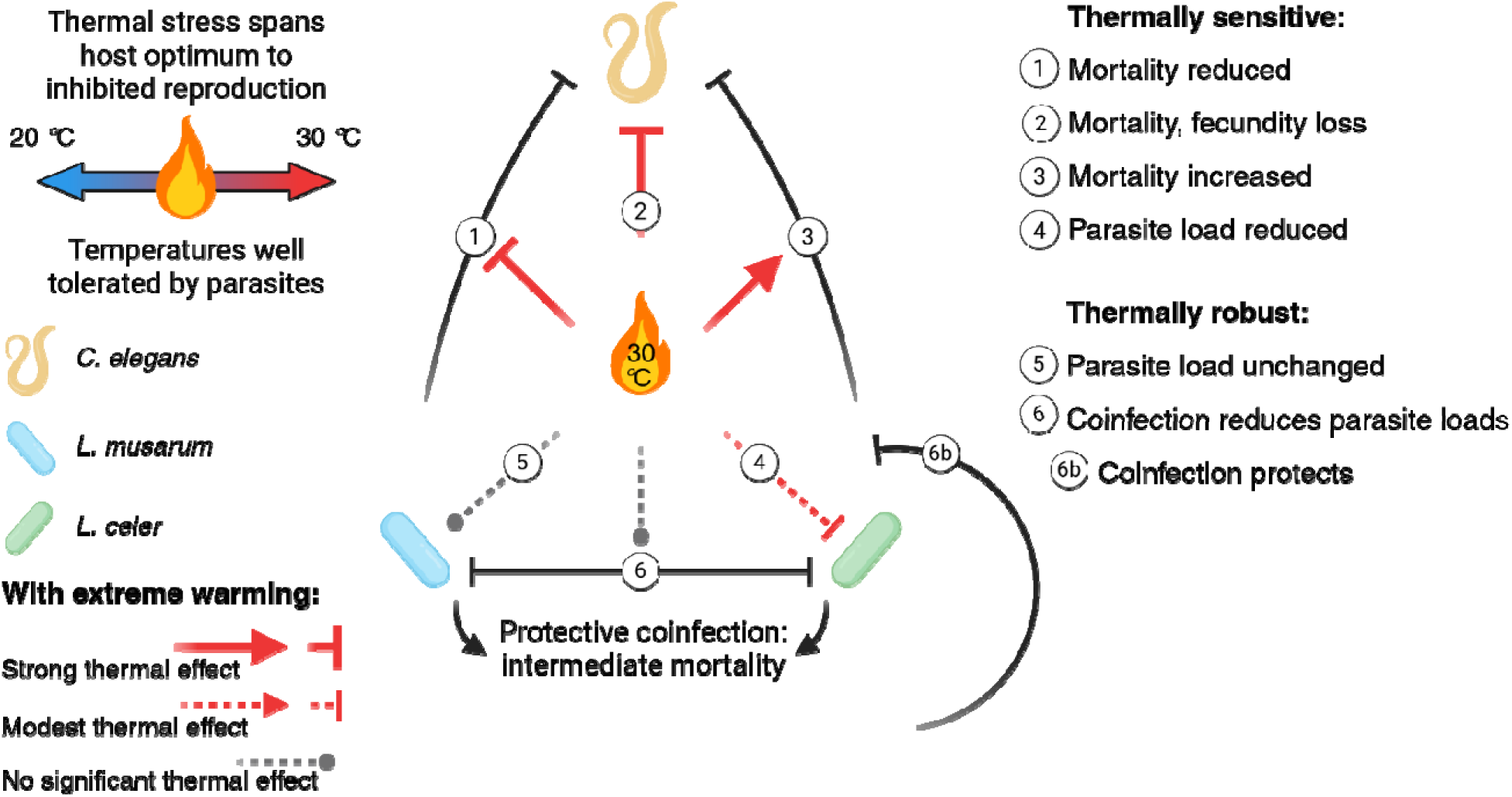
Extreme heating revealed thermally sensitive and thermally robust interactions that reflect host and parasite thermal tolerance. When hosts experience extreme thermal stress (30 °C), disease outcomes change significantly, but with contrasting parasite-specific effects. Parasite-parasite coinfection interactions continued to be competitive and provide protection. Parasite coinfection interactions thus dampened the possible effect of thermal stress on disease. Arrows indicate a causative link (e.g., coinfection producing intermediate mortality) or an increased effect (e.g., warming increased mortality from *L. celer*). Bars show harm (e.g. coinfection reduces parasite loads) or a reduced effect (e.g., warming reduced mortality from *L. musarum*). Key effects are labeled with numbers, but do not necessarily reflect a sequence of events.

Across all temperatures, coinfection resulted in host protection (Figure 3). *Leucobacter celer* coinfection protected hosts against *L. musarum* during 20 °C, 25 °C, and heatwave treatments (Figure 3A, Table S3, Supporting data S1). At 30 °C, where LC infection was most lethal, coinfection by *L. musarum* was also protective, albeit to a lesser degree (Figure 3A, Figure 4, Table S3, Supporting data S1). This smaller protective effect at 30 °C corresponded to a 34% mortality reduction, compared to 71-77% reductions conferred by *L. celer* at other temperatures (Figure 3A). We paired our measurements of host mortality with a sub-lethal aspect of host fitness, host fecundity in individuals that survived infection. Coinfection continued to partially protect hosts, buffering fecundity to intermediate levels. The fecundity assay methods and results are detailed in Supporting document S1.

All infections caused less mortality during heatwave than would be predicted by their effect at 20 °C (Figure 3, Table S3). Acute heatwave exposure to 35 °C drove the highest levels of mortality in uninfected *C. elegans* (13.6% median mortality). This increased “baseline” mortality accounted for most of the 25.6% median mortality observed during LM-heatwave treatments. An LM by heatwave interaction significantly reduced the mortality effect of LM infections compared to 20 °C LM (Table S3). Infections including *L. celer* (LC or coinfection) were avirulent, or, possibly, protected against thermal stress (Table S3, Supporting data S1). Although protection against thermal stress is biologically plausible, given that that the median observed control-heatwave mortality lies much closer to LC or coinfection than the model estimate (Figure 3A), we leave this possible effect for future validation.

LC infections with higher loads did not increase mortality, whereas LM infections likely did across temperatures (Figure 3B, Supporting document S3). Coinfection only had a temperature specific effect, at 30 °C (Figure 3B, Supporting document S3). At 30 °C, total coinfection loads appeared to have a reducing effect on host mortality (Figure 3B) – however, underlying this observation was that the two species had opposing impacts on mortality (Figure 3, Supporting document S3). Modeling coinfected host mortality by loads of each species separately revealed load-temperature interactions at 30 °C; a positive relationship between host mortality and *L. celer* loads (scaled per-CFU effect = 0.93, 95CI = 0.39, 1.47), was counteracted by a negative relationship with *L. musarum* loads that reduced host mortality (effect = - 1.84, 95CI = −2.71, −0.96) (see Supporting document S3 for a full consideration of the mortality by load models).

### Coinfections continued to buffer disease outcomes in hosts with altered susceptibility

We tested if protection conferred by coinfection persisted despite altered host-parasite interaction in contexts other than temperature alone. We used *srf-2* mutant *C. elegans* that have inversed susceptibilities to the *Leucobacter* parasites (Hodgkin *et al*. 2013; Li *et al*. 2025), thus directly changing host-parasite interactions (Figure 5). We created a GLMM of *srf-2* host mortality, in the same form as that used for N2 host data: “Mortality ∼ Temperature*Infection + (1|Batch).” This *srf-2* model had both conditional and marginal Bayes R^2^ of 0.97. We found coinfection continued to protect against the most severe possible effects of temperature on host mortality in this mutant line. However, with sustained high temperature, the strength of that protective coinfection was greatly reduced (Figure 5; Table S4).

**Figure 5.**
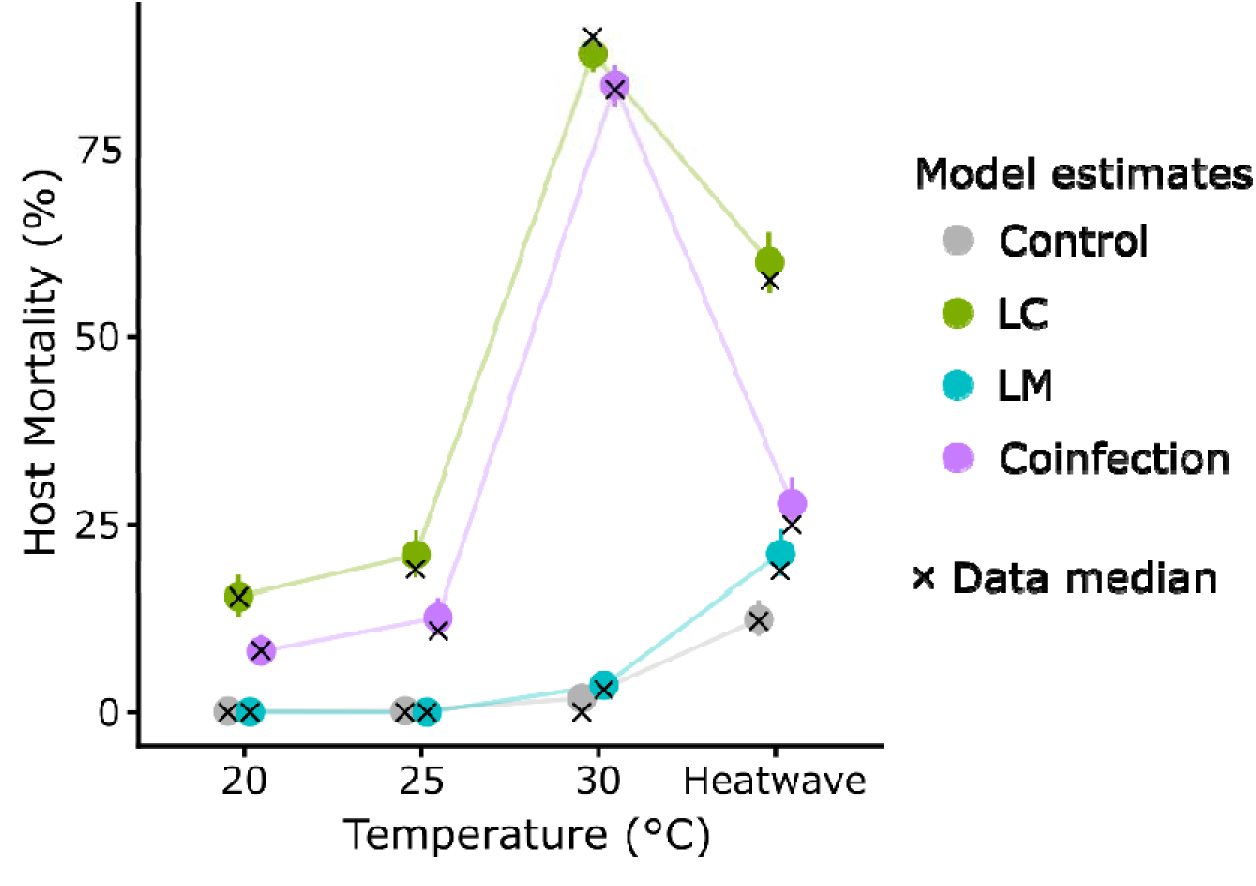
Parasite interactions continued to produce intermediate mortality in genetically susceptible hosts, except with the temperature that drove the highest virulence. Model estimates (points) and 95CI (bars) show LM (*L. musarum* alone) and LC (*L. celer* alone) have inversed mortality outcomes compared to N2 genotype hosts at lower temperatures and exacerbated LC mortality at higher temperatures (Figure 3). However, parasite competition was still capable of protecting *srf-2* mutant hosts, leading to intermediate mortality. This protection was notably less effective with combined thermal stress and genetic susceptibility (LC, *srf-2*, and 30 °C combination). Single infection LC treatments produced mass mortality at 30 °C (median 90% mortality). Median values of the raw data (Figure S7) are given by crosses. Points are connected by lines to show trends.

Coinfecting parasites typically produced intermediate mortality in the *srf-2* host genotype (Figure 5), as was found for N2 hosts (Figure 3). However, combining *srf-2* host genetic susceptibility to *L. celer* and extreme thermal stress (30 °C) severely eroded the protective effect of coinfection. Coinfection interactions benefited *srf-2* hosts with 43-57% mortality reductions at 20 °C, 25 °C, and heatwave (Figure 5). At 30 °C, coinfection was no longer clearly protective (estimate 95CI for coinfection-30 °C = 80.6, 86.2 and LC-30 °C = 85.3, 89.9) (Figure 5, Table S4). We considered the implications of temperature-sensitive protection for short-term population dynamics in Figure S9. There, we highlight the theoretical possibility for protection to increase transmission of parasites and increase net mortality if the environment later undergoes warming.

To test if mass mortality during LC infection was due to a specific genotype by environment interaction, we created an additional GLMM. This GLMM can be summarized as “Mortality ∼ Temperature*Genotype + (1|Batch)” and only used data from LC treatment, for both N2 and *srf-2* hosts (Table S5). This model had a conditional Bayes R^2^ of 0.95 and a marginal of 0.94. Both the susceptible *srf-2* genotype and 30 °C temperature had strong positive effect sizes that reflected increased host mortality (Table S5). These separate, additive genotype and environment effects were sufficient to explain mass host mortality, without a clear genotype by environment interaction (Table S5).

### Temperature modifies single-parasite virulence across protective coinfection scenarios

We used mathematical modeling to understand how contrasting disease outcomes across conditions, such as those we observed with temperature, can occur more generally. We modeled disease scenarios exemplified by three characteristics of our experimental system, (*i*) non-linear, temperature-dependent mortality, (*ii*) environmental transmission of parasites, and (*iii*) coinfection interactions leading to intermediate mortality. The number of infected individuals (transmission, β) and the rate at which they die (virulence, α) could vary to give population-level mortality (Figure 1). Following our empirical data, we constrained these models by requiring coinfections to produce intermediate mortality. As the exact mechanism of protection is unclear, we considered two possible routes to lower mortality, either reduced transmission (exclusion) or reduced virulence (attenuation). In essence, these modes of protection could represent mechanisms preventing colonization of hosts from the environment (exclusion) or limitation of the competitor’s growth within-hosts and/or conferral of tolerance (attenuation). We specified temperature-dependent levels of single-infection mortality based on our experiments (mean mortality per temperature, per *Leucobacter* species) and made a simplifying assumption that the strength of protection was independent of temperature.

Coinfection highlighted a consistently important role for virulence in moving between temperature-dependent mortality levels (Figure 6, Supporting document S2). When parasites excluded the competitor species, virulence was necessary and sufficient to traverse the theoretical mortality landscape based on both *L. celer* (Figure 6A) and *L. musarum* (Figure 6B) host mortality levels. When coinfection attenuated disease, mortality due to *L. celer* was also explained best by virulence (Figure 6C).

**Figure 6.**
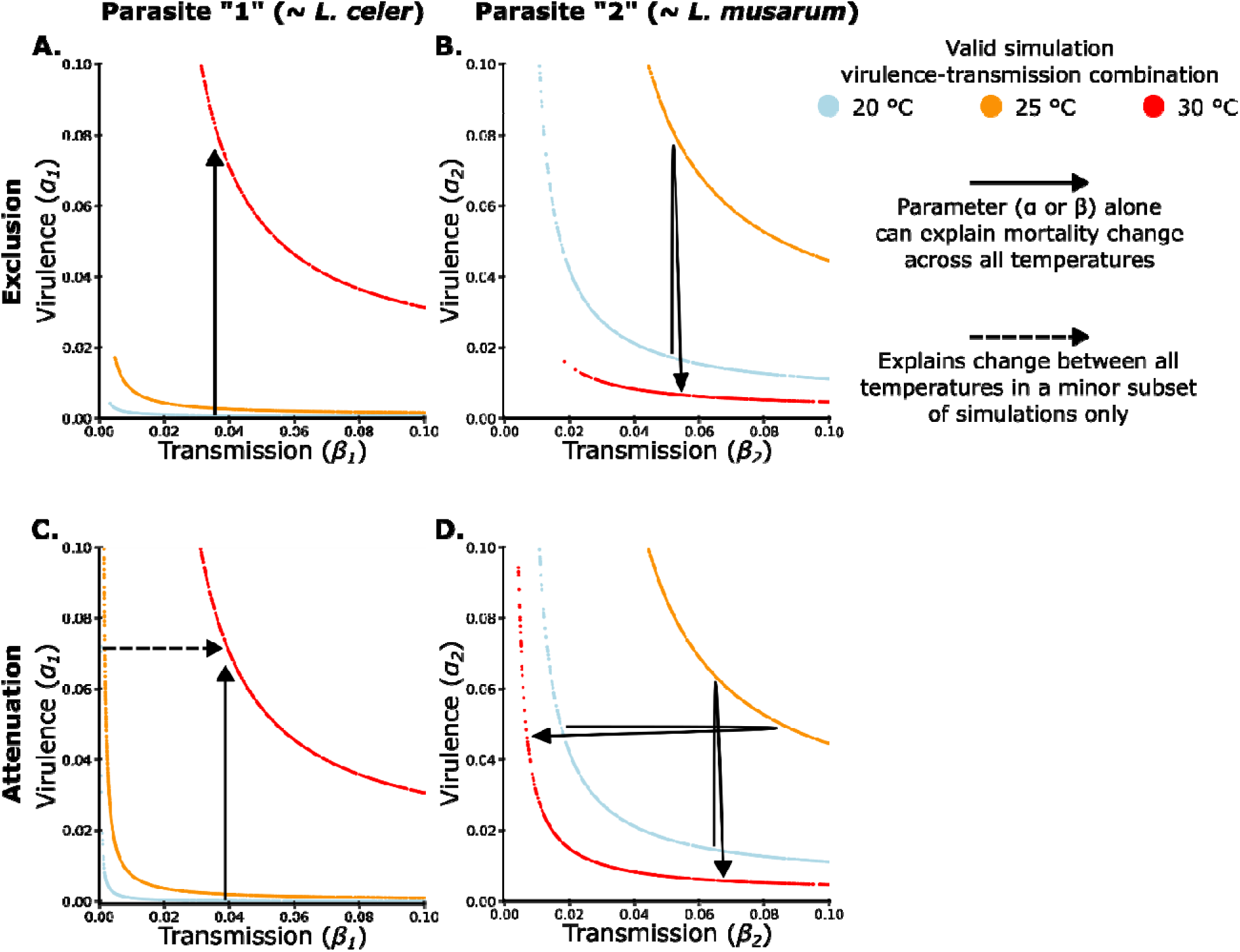
Virulence was always sufficient to shift single-infection mortality across temperatures when parasites competed. Protective parasite competition, which limited population-level host mortality to intermediate levels, constrained the ways by which changes in transmission or virulence could traverse the theoretical mortality landscape (Figure 1). Virulence was always sufficient to explain all shifts due to temperature, whether parasite competition excluded invasion of a second parasite (**A.**, **B.**) or attenuated virulence within coinfected individuals (**C.**, **D.**). The contours (each point is a simulation) for three levels of single infection mortality were constructed using mean mortality values taken from our experimental data, as a representative case. We applied our general model to these experimental mortality values found with *L. celer* (as “parasite 1”) and *L. musarum* (as “parasite 2”). Simulations were filtered such that cases in which parasite competition produced intermediate mortality were plotted, producing the truncated single-infection mortality curves displayed. The black arrows denote how virulence or transmission changes can most simply traverse the mortality landscape from 20 to 30 °C. Possibilities exist for combined transmission and virulence changes, but we show the simplest cases, requiring only one to change. The dashed arrow in (**C.**) indicates transmission could explain all mortality changes, but only for very limited parameter values (4/1,000 simulations at 20 °C).

Transmission only plausibly drove all observed mortality changes for *L. musarum* infections in the protection by attenuation case (Figure 6D).

## DISCUSSION

Coinfection and temperatures can independently contribute to variation in disease outcomes in the wild (Hoarau *et al*. 2020; Paull & Johnson 2014; Pedersen & Fenton 2007). We found that temperature had disparate impacts on host mortality and loads of two parasite species during single-infections. When parasites competed in coinfection, however, virulence was buffered from otherwise dramatic changes in host mortality driven by the combination of heating and disease.

In “thermal mismatches” when the host experiences more thermal stress than its parasites, disease can become exacerbated (Cohen *et al*. 2017, 2019, 2020; Sauer *et al*. 2020). However, we observed distinct virulence outcomes that deviated from the simplest predictions based on thermal performance alone. Sustained high temperature (30 °C) shifted a benign infection to highly virulent, and vice versa, revealing contrasting host-parasite-environment relationships. Environmental growth of the parasites predicted a thermal advantage over hosts at 30 °C, but the within-host parasite loads were either reduced or unchanged relative to the host optimum temperature. The within-host environment, at this physiological extreme, appears to have produced deviations from expectations based on thermal mismatch or metabolic theory. Although, we found some congruence of load and virulence at intermediate temperatures (20 and 25 °C) for *L. musarum* infections (as in Hector *et al*. 2025). The specific ecology of parasite species (Slowik *et al*. 2024), variation in host immunity-parasite interactions (Mignatti *et al*. 2016), and mode of transmission or biological scale (i.e., individuals or populations) (Kirk *et al*. 2022) have all been found to add complexity to temperature-disease interactions.

We found thermal stress introduced such complexity when crossing from physiological to extreme temperatures for the host. Thermal stress and immunity are linked at the molecular level in *C. elegans* (Feder & Hofmann 1999; Hsu *et al*. 2003; Singh & Aballay 2006; Sørensen *et al*. 2003) and, at 30 °C, *C. elegans* experiences reproduction-halting levels of stress (Begasse *et al*. 2015; Gouvêa *et al*. 2015; Petrella 2014; Plagens *et al*. 2021). Shifts in fundamental reproduction-defense host fitness tradeoffs (i.e., dynamic energy budgets) possibly explained the divergent disease outcomes we observed. For example, castrating parasites can induce hypertrophy (somatic investment) in their hosts (Hall *et al*. 2007) and germline-free *C. elegans* better survive bacterial infections (Alper *et al*. 2009; Miyata *et al*. 2008; Tekippe & Aballay 2010). Host populations at reproduction-limiting temperatures may also be poised to collapse below the required “threshold density” to maintain parasite populations long-term, where the thermal mismatch hypothesis predictions are most robust (Lloyd-Smith *et al*. 2005; Rohr & Cohen 2020). Although firm thresholds may only be rarely observed in the wild (Lloyd-Smith *et al*. 2005), parasites may have limited opportunity to adapt to highly stressed hosts that historically have been sparsely or transiently present. Variation in host and parasite responses to increasingly erratic climate conditions complicate forecasting disease outcomes in emerging disease scenarios (Bull & Ebert 2008; Ekroth *et al*. 2021; Vasseur *et al*. 2014).

Differences in virulence and transmission produce variation in disease outcomes. Although virulence and transmission can be linked by parasite fitness tradeoffs (Anderson & May 1982; De Roode *et al*. 2008), their relationship can also be complex and dependent on other factors (Alizon & Michalakis 2015; Handel & Rohani 2015; Silva *et al*. 2025). Virulence, rather than transmission, most robustly explained temperature-dependent mortality in our mathematical models. In contrast, other systems (e.g., tested without protective coinfections) have found temperature changes during single infections to significantly alter transmission (Ismail *et al*. 2024; Suh *et al*. 2024). In light of these findings, we equate changes in mortality in our experiments as changes in virulence. Future experiments would be required to reveal the molecular mechanism by which temperature modifies virulence.

Distinguishing virulence and transmission effects across temperatures is a key component of evaluating disease risks and possible interventions (Nguyen *et al*. 2021; Paull & Johnson 2014). Furthermore, modeling scenarios where protective coinfection is thermally sensitive (e.g., *srf-2* mutants at 30 °C) could reveal different infection dynamics, such as those that lead to mass mortality.

Coinfected hosts died less across temperatures than hosts singly-infected with the most virulent disease. Competition reduced loads of both parasite species, effectively conferring infection resistance to hosts and likely underpinning protection. Although virulence-load links on a per-bacterium basis within samples had mixed results, the overall trend of coinfection reducing both host mortality and loads of both parasites was clear. Coinfection interactions did not substantially shift while temperature and host genotype led to major changes between host and parasite. Transcriptomic work in this system has hypothesized that protective coinfection may operate by exploitative competition between parasites for “public goods”, operating in parallel to endogenous host defenses (Will *et al*. 2025). This parasite-parasite competition may alleviate immunity-reproduction tradeoffs in the host resource budget (Will *et al*. 2025), which could offset temperature-dependent changes in host-parasite interactions. Although temperature can alter the outcomes of microbial antagonisms (Dal Bello & Abreu 2024; Li *et al*. 2022), we found that these *Leucobacter* parasites were thermally matched and competition reduced loads of both across temperatures. We additionally tested mutant hosts, with inversed susceptibilities to the *Leucobacter* species (Hodgkin *et al*. 2013; Li *et al*. 2025; Loer *et al*. 2015; O’Rourke *et al*. 2023; Parsons *et al*. 2014), and found coinfection remained protective. Parasite-parasite conflict can thus affect hosts across contexts – dampening the impacts of either environment (e.g., temperature) or host biology (e.g., genotype) on variation in virulence. We nevertheless demonstrated that combined environmental and host condition factors can nearly wipe out host populations, whether singly- or coinfected. This finding supports observations in the wild for mass mortality events being commonly driven by infectious disease, with additional stressors enhancing virulence (Carella *et al*. 2019; Fey *et al*. 2015; Sanderson & Alexander 2020). Further research could explore conditions where the parasites themselves are thermally mismatched and if different modes of coinfection interactions (e.g., exploitative competition for host nutrients, spiteful molecular warfare, cooperation, or sequential exposure) (Ramesh & Hall 2025) affect disease in novel ways. Parasite interactions across large taxonomic divides, utilizing distinct host-resources, or interacting at a distance between host-compartments (Erazo *et al*. 2025; Graham 2008; Lello *et al*. 2018), could capture different modes of competition than we observed between the congener species we tested.

In a warming world with parasites spreading to areas undergoing rapid heating and environmental change, temperate host species (and their local parasites) may be challenged by novel parasite threats from warmer regions (Carlson *et al*. 2022). Coinfecting parasites may spread in tandem (Kafle *et al*. 2020; Oakgrove *et al*. 2014). Protective coinfection could mitigate the worst possible outcomes from emerging infectious disease. As coinfections are widespread (Telfer *et al*. 2008), predictions based purely on single infection thermal mismatches or metabolic first principles may misestimate the impacts on animal health when not accounting for inter-parasite interactions. Protective coinfections in particular may reduce host mortality levels in the short term but may facilitate the rapid transmission of both parasites across time. These widespread infections could be a latent source of severe disease, emerging when protection breaks down.

## Acknowledgements

KCK was funded by a European Research Council grant (COEVOPRO #802242) and Natural Environment Research Council (NE/X000540/1). The *srf-2* mutant line was generously gifted by Johnathan Hodgkin. We would like to thank Mathilda J. Whittle for helpful discussions on mathematically representing the loss of coinfection protection and Michael Blazanin for comments on an early version of the manuscript. Figure 4 was created with Biorender.com.

## Authorship

IW and KCK designed the experiments. IW and TEH collected the data. CAS developed the mathematical models. IW conducted the statistical analyses. IW, CAS, TEH, and KCK wrote the manuscript.

## Data availability

The accompanying code for the mathematical mode can be found at: https://doi.org/10.5281/zenodo.17897997

## Supporting information

**Supporting data S1.** Excel file with posterior estimates for the N2 host mortality model across reference levels.

**Table S1.** A model of *L. celer* loads indicated that bacterial abundance was reduced both by coinfection and constant high temperature (30 °C).

**Table S2.** A model of *L. musarum* loads indicated that bacterial abundance was maximal at 25 °C and suppressed by coinfection except during heatwave.

**Table S3.** A model of host mortality indicated clear temperature, infection, and interactive effects.

**Table S4.** A model of *srf-2* host mortality found that coinfection interactions continued to protect hosts, even has temperature drove large changes in mortality.

**Table S5.** A model of LC host mortality indicated clear genotype and temperature effects, but their interaction did not drive mass mortality.

**Figure S1.** Using a fully factorial design, we analyzed up to seven batches of N2 genotype infection assays to produce the primary dataset.

**Figure S2.** Nematode washing removed the vast majority of bacteria not directly associated with the host.

**Figure S3.** Parasite coinfection competition mediated by the host appears to be stronger than environmental competition alone.

**Figure S4.** Maximum growth rate observed in single cultures of *Leucobacter* grown *in vitro* at different temperatures.

**Figure S5.** Parasite fitness (CFU load) was generally reduced by coinfection, with secondary effects of temperature.

**Figure S6.** Coinfecting parasite interactions modified temperature-dependent host mortality.

**Figure S7.** Parasite interactions persisted to give intermediate mortality, despite major changes in host condition due to genetic susceptibility.

**Figure S8.** Bacterial competition is typically mutually harmful.

**Figure S9.** Increased survivorship due to protection could lead to greater parasite transmission and downstream increases in deaths.

**SUPPORTING document S1.** Host fecundity is primarily shaped by temperature and secondarily by infection.

**SUPPORTING document S2.** Mathematical models.

**Supporting document S3.** Severe warming may link coinfecting parasite loads with virulence and protection.

