## Supplemental Information for "Coinfections dampen the effects of temperature on host-parasite interactions"

**Supporting information:**

**Supporting data S1.** Excel file with posterior estimates for the N2 host mortality model across reference levels.

**Table S1.** A model of *L. celer* loads indicated that bacterial abundance was reduced both by coinfection and constant high temperature (30 °C).

**Table S2.** A model of *L. musarum* loads indicated that bacterial abundance was maximal at 25 °C and suppressed by coinfection except during heatwave.

**Table S3.** A model of host mortality indicated clear temperature, infection, and interactive effects.

**Table S4.** A model of *srf-2* host mortality found that coinfection interactions continued to protect hosts, even has temperature drove large changes in mortality.

**Table S5.** A model of LC host mortality indicated clear genotype and temperature effects, but their interaction did not drive mass mortality.

**Figure S1.** Using a fully factorial design, we analyzed up to seven batches of N2 genotype infection assays to produce the primary dataset.

**Figure S2.** Nematode washing removed the vast majority of bacteria not directly associated with the host.

**Figure S3.** Parasite coinfection competition mediated by the host appears to be stronger than environmental competition alone.

**Figure S4.** Maximum growth rate observed in single cultures of *Leucobacter* grown *in vitro* at different temperatures.

**Figure S5.** Parasite fitness (CFU load) was generally reduced by coinfection, with secondary effects of temperature.

**Figure S6.** Coinfecting parasite interactions modified temperature-dependent host mortality.

**Figure S7.** Parasite interactions persisted to give intermediate mortality, despite major changes in host condition due to genetic susceptibility.

**Figure S8.** Bacterial competition is typically mutually harmful.

**Figure S9.** Increased survivorship due to protection could lead to greater parasite transmission and downstream increases in deaths.

**Supporting document S1.** Host fecundity is primarily shaped by temperature and secondarily by infection.

**Supporting document S2.** Mathematical models.

**Supporting document S3.** Severe warming may link coinfecting parasite loads with virulence and protection.

**Supporting data S1**

An excel table showing outputs of brms function hypothesis() to obtain posterior 95CI and estimates for main effects of temperature or infection with different reference levels than “control:20 °C” as primarily reported in the main text.

**Table S1.** **A model of *L. celer* loads indicated that bacterial abundance was reduced both by coinfection and constant high temperature (30 °C).** This frequentist GLMM (negative binomial) can be summarized as “CFUs ~ Temperature*Infection + (1|Batch)”. The model had a Nakagawa’s pseudo-R^2^ of 0.601 (conditional) and 0.441 (marginal). Batch was a five-level random effect. Variable abbreviations: “Temp.HW” is heatwave, “Inf.Con” is control, and “Inf.Coinf” is coinfection. (**A**) Interaction terms are given by a colon. Significant p-values (≤ 0.05) are in bold. (**B**) The second part of the table gives the treatment level model estimates on the mortality probability response scale. These treatment levels consist of a temperature and infection pair, given by a colon. Uncertainty is given by frequentist 95% confidence intervals, not the credible intervals.

| **(A) Independent variable** | **Coefficient** | **Error** | **Test statistic (z)** | **p-value** |
| --- | --- | --- | --- | --- |
| Intercept (Temp.20 : Inf.LC) | 11.11 | 0.27 | 41.84 | < 2e-16 |
| Temp.25 | -0.41 | 0.29 | -1.40 | 0.161 |
| **Temp.30** | **-0.85** | **0.29** | **-2.89** | **0.004** |
| Temp.HW | -0.38 | 0.29 | -1.30 | 0.194 |
| **Inf.Coinf** | **-1.32** | **0.29** | **-4.48** | **7.51e-06** |
| Temp.25 : Inf.Coinf | 0.45 | 0.41 | 1.10 | 0.272 |
| Temp.30 : Inf.Coinf | 0.10 | 0.41 | 0.24 | 0.812 |
| Temp.HW : Inf.Coinf | 0.58 | 0.43 | 1.34 | 0.179 |
| Random effect (Batch): 0.140 variance, 0.375 standard deviation | | | | |
| **(B) Treatment** | **Estimate** | **Error** | **95% Confidence, low** | **95% Confidence, high** |
| Temp.20 : Inf.LC | 66511.66 | 0.2654 | 39533.63 | 111899.70 |
| Temp.25 : Inf.LC | 44278.30 | 0.2645 | 26368.03 | 74353.97 |
| Temp.30 : Inf.LC | 28534.88 | 0.2686 | 16856.38 | 48304.52 |
| Temp.HW : Inf.LC | 45474.19 | 0.2659 | 27003.54 | 76578.93 |
| Temp.20 :Inf.Coinf | 17820.61 | 0.2667 | 10565.86 | 30056.64 |
| Temp.25 : Inf.Coinf | 18596.96 | 0.2667 | 11027.17 | 31363.16 |
| Temp.30 : Inf.Coinf | 8438.83 | 0.2675 | 4995.459 | 14255.72 |
| Temp.HW : Inf.Coinf | 21702.63 | 0.2729 | 12712.26 | 37051.18 |

**Table S3.** **A model of *L. musarum* loads indicated that bacterial abundance was maximal at 25 °C and suppressed by coinfection except during heatwave.** This frequentist GLMM (negative binomial) can be summarized as “CFUs ~ Temperature*Infection + (1|Batch)”. The model had a Nakagawa’s pseudo-R^2^ of 0.620 (conditional) and 0.479 (marginal). Batch was a five-level random effect. Abbreviations are as described in Table S1. Significant p-values (≤ 0.05) are in bold. (**A**) Interaction terms are given by a colon. Significant p-values (≤ 0.05) are in bold. (**B**) The second part of the table gives the treatment level model estimates on the mortality probability response scale. These treatment levels consist of a temperature and infection pair, given by a colon. Uncertainty is given by frequentist 95% confidence intervals, not the credible intervals.

| **(A) Independent variable** | **Coefficient** | **Error** | **Test statistic (z)** | **p-value** |
| --- | --- | --- | --- | --- |
| Intercept (Temp.20 : Inf.LM) | 10.77 | 0.24 | 45.49 | 2e-16 |
| **Temp.25** | **0.93** | **0.26** | **3.54** | **4.03e-04** |
| Temp.30 | -0.11 | 0.26 | -0.43 | 0.665 |
| Temp.HW | 0.13 | 0.26 | 0.51 | 0.607 |
| **Inf.Coinf** | **-1.07** | **0.26** | **-4.10** | **4.17e-05** |
| Temp.25 : Inf.Coinf | 0.26 | 0.37 | 0.70 | 0.482 |
| Temp.30 : Inf.Coinf | 0.34 | 0.37 | 0.92 | 0.355 |
| **Temp.HW : Inf.Coinf** | **1.35** | **0.37** | **3.63** | **2.88e-04** |
| Random effect (Batch): 0.109 variance, 0.330 standard deviation | | | | |
| **(B) Treatment** | **Estimate** | **Error** | **95% Confidence, low** | **95% Confidence, high** |
| Temp.20 : Inf.LC | 47333.01 | 0.2366 | 29767.18 | 75264.56 |
| Temp.25 : Inf.LC | 120507.73 | 0.2388 | 75471.35 | 192418.90 |
| Temp.30 : Inf.LC | 42253.75 | 0.2374 | 26531.51 | 67292.81 |
| Temp.HW : Inf.LC | 54147.32 | 0.2368 | 34044.85 | 86119.69 |
| Temp.20 :Inf.Coinf | 16185.57 | 0.2375 | 10161.49 | 25780.91 |
| Temp.25 : Inf.Coinf | 53553.98 | 0.2377 | 33612.21 | 85326.99 |
| Temp.30 : Inf.Coinf | 20382.56 | 0.2411 | 12707.19 | 32694.00 |
| Temp.HW : Inf.Coinf | 71582.61 | 0.2398 | 44735.81 | 114540.7 |

**Table S3.** **A model of host mortality indicated clear temperature, infection, and interactive effects.** This Bayesian logistic regression can be summarized as “Mortality ~ Temperature*Infection + (1|Batch)”. The model had a conditional (including random effects) Bayes R^2^ of 0.75 and a marginal (fixed effects only) of 0.54 (standard error of 0.01). Batch was a seven-level random effect. Column “Error” is the standard error. (**A**) Interaction terms are given by a colon. 95CIs that do not include zero are shown in bold. The coefficient values are logits. (**B**) The second part of the table gives the treatment level model estimates on the mortality probability response scale. These treatment levels consist of a temperature and infection pair, given by a colon. Estimates with non-overlapping 95CI intervals are considered to have clearly different levels of mortality.

| **(A) Independent variable** | **Coefficient** | **Error** | **95CI, low** | **95CI, high** |
| --- | --- | --- | --- | --- |
| Intercept (Temp.20 : Inf.Con) | -6.65 | 0.53 | -7.76 | -5.68 |
| Temp.25 | 0.61 | 0.65 | -0.67 | 1.89 |
| **Temp.30** | **2.03** | **0.54** | **1.02** | **3.14** |
| **Temp.HW** | **5.45** | **0.50** | **4.54** | **6.50** |
| **Inf.LC** | **1.22** | **0.58** | **0.10** | **2.40** |
| **Inf.LM** | **4.91** | **0.50** | **4.00** | **5.97** |
| **Inf.Coinf** | **3.64** | **0.51** | **2.71** | **4.70** |
| Temp.25 : Inf.LC | 0.37 | 0.74 | -1.07 | 1.84 |
| **Temp.30 : Inf.LC** | **2.25** | **0.62** | **1.01** | **3.45** |
| **Temp.HW : Inf.LC** | **-2.07** | **0.59** | **-3.25** | **-0.94** |
| Temp.25 : Inf.LM | 0.51 | 0.66 | -0.77 | 1.79 |
| **Temp.30 : Inf.LM** | **-3.25** | **0.56** | **-4.38** | **-2.20** |
| **Temp.HW : Inf.LM** | **-4.60** | **0.51** | **-5.66** | **-3.68** |
| Temp.25 : Inf.Coinf | 0.09 | 0.66 | -1.21 | 1.39 |
| Temp.30 : Inf.Coinf | -0.74 | 0.56 | -1.87 | 0.30 |
| **Temp.HW : Inf.Coinf** | **-4.62** | **0.52** | **-5.71** | **-3.67** |
| Random effect (Batch) | 0.46 | 0.19 | 0.24 | 0.93 |
| **(B) Treatment** | **Estimate** | **Error** | **95CI, low** | **95CI, high** |
| Temp.20 : Inf.Con | 1.14E-03 | 5.47E-04 | 3.93E-04 | 2.75E-03 |
| Temp.25 : Inf.Con | 2.10E-03 | 9.55E-04 | 7.32E-04 | 4.74E-03 |
| Temp.30 : Inf.Con | 8.38E-03 | 2.00E-03 | 5.04E-03 | 1.30E-02 |
| Temp.HW : Inf.Con | 2.04E-01 | 1.36E-02 | 1.78E-01 | 2.31E-01 |
| Temp.20 : Inf.LC | 3.80E-03 | 1.30E-03 | 1.78E-03 | 7.06E-03 |
| Temp.20 : Inf.LM | 1.30E-01 | 1.06E-02 | 1.10E-01 | 1.52E-01 |
| Temp.20 : Inf.Coinf | 4.02E-02 | 5.12E-03 | 3.11E-02 | 5.13E-02 |
| Temp.25 : Inf.LC | 1.00E-02 | 2.23E-03 | 6.24E-03 | 1.51E-02 |
| Temp.30 : Inf.LC | 2.13E-01 | 1.48E-02 | 1.86E-01 | 2.44E-01 |
| Temp.HW : Inf.LC | 9.89E-02 | 8.95E-03 | 8.26E-02 | 1.18E-01 |
| Temp.25 : Inf.LM | 3.14E-01 | 1.74E-02 | 2.80E-01 | 3.49E-01 |
| Temp.30 : Inf.LM | 4.24E-02 | 5.39E-03 | 3.28E-02 | 5.40E-02 |
| Temp.HW : Inf.LM | 2.58E-01 | 1.60E-02 | 2.28E-01 | 2.91E-01 |
| Temp.25 : Inf.Coinf | 7.78E-02 | 7.91E-03 | 6.35E-02 | 9.46E-02 |
| Temp.30 : Inf.Coinf | 1.32E-01 | 1.10E-02 | 1.12E-01 | 1.55E-01 |
| Temp.HW : Inf.Coinf | 8.70E-02 | 8.56E-03 | 7.12E-02 | 1.05E-01 |

**Table S4. A model of *srf-2* host mortality found that coinfection interactions continued to protect hosts, even as temperature drove large changes in mortality.** This Bayesian logistic regression model can be summarized as “Mortality ~ Temperature*Infection + (1|batch)”. The model had both a conditional and marginal Bayes R^2^ of 0.97 (standard error of 0.001). We included a four-level random effect of “Batch”. This model was for *srf-2* mutant *C. elegans* only. Coinfection remained protective across temperatures but was less effective at 30 °C. Abbreviations are as described in Table S1. (**A**) Interaction terms are given by a colon. 95CIs that do not include zero are shown in bold. The coefficient values are logits. (**B**) The second part of the table gives the treatment level model estimates on the mortality probability response scale. These treatment levels consist of a temperature and infection pair, given by a colon.

| **(A) Independent variable** | **Coefficient** | **Error** | **95CI, low** | **95CI, high** |
| --- | --- | --- | --- | --- |
| Intercept (Temp.20 : Inf.Con) | -6.77 | 0.65 | -8.11 | -5.58 |
| Temp.25 | 0.26 | 0.85 | -1.46 | 1.87 |
| **Temp.30** | **2.71** | **0.64** | **1.52** | **4.03** |
| **Temp.HW** | **4.73** | **0.62** | **3.61** | **6.01** |
| **Inf.LC** | **4.98** | **0.62** | **3.86** | **6.26** |
| Inf.LM | -1.13 | 1.13 | -3.54 | 0.93 |
| **Inf.Coinf** | **4.26** | **0.62** | **3.13** | **5.55** |
| Temp.25 : Inf.LC | 0.12 | 0.86 | -1.5 | 1.86 |
| Temp.30 : Inf.LC | 0.97 | 0.65 | -0.37 | 2.18 |
| **Temp.HW : Inf.LC** | **-2.62** | **0.62** | **-3.91** | **-1.47** |
| Temp.25 : Inf.LM | -3.18 | 2.36 | -8.18 | 0.99 |
| Temp.30 : Inf.LM | 1.81 | 1.16 | -0.3 | 4.26 |
| Temp.HW : Inf.LM | 1.77 | 1.14 | -0.3 | 4.18 |
| Temp.25 : Inf.Coinf | 0.23 | 0.86 | -1.4 | 1.98 |
| **Temp.30 : Inf.Coinf** | **1.34** | **0.66** | **4e-3** | **2.56** |
| **Temp.HW : Inf.Coinf** | **-3.26** | **0.63** | **-4.56** | **-2.11** |
| Random effect (Batch) | 0.36 | 0.31 | 0.11 | 1.17 |
| **(B) Treatment** | **Estimate** | **Error** | **95CI, low** | **95CI, high** |
| Temp.20 : Inf.Con | 1.28E-03 | 7.58E-04 | 3.47E-04 | 3.77E-03 |
| Temp.25 : Inf.Con | 1.71E-03 | 1.10E-03 | 3.53E-04 | 5.17E-03 |
| Temp.30 : Inf.Con | 1.85E-02 | 4.37E-03 | 1.13E-02 | 2.86E-02 |
| Temp.HW : Inf.Con | 1.24E-01 | 1.22E-02 | 1.01E-01 | 1.49E-01 |
| Temp.20 : Inf.LC | 1.53E-01 | 1.39E-02 | 1.28E-01 | 1.82E-01 |
| Temp.20 : Inf.LM | 4.45E-04 | 4.34E-04 | 3.50E-05 | 2.63E-03 |
| Temp.20 : Inf.Coinf | 8.13E-02 | 1.02E-02 | 6.27E-02 | 1.03E-01 |
| Temp.25 : Inf.LC | 2.10E-01 | 1.63E-02 | 1.79E-01 | 2.43E-01 |
| Temp.30 : Inf.LC | 8.78E-01 | 1.16E-02 | 8.53E-01 | 8.99E-01 |
| Temp.HW : Inf.LC | 6.00E-01 | 2.07E-02 | 5.58E-01 | 6.39E-01 |
| Temp.25 : Inf.LM | 2.82E-05 | 4.05E-05 | 1.01E-07 | 1.09E-03 |
| Temp.30 : Inf.LM | 3.60E-02 | 6.33E-03 | 2.50E-02 | 5.01E-02 |
| Temp.HW : Inf.LM | 2.11E-01 | 1.59E-02 | 1.81E-01 | 2.43E-01 |
| Temp.25 : Inf.Coinf | 1.26E-01 | 1.26E-02 | 1.03E-01 | 1.52E-01 |
| Temp.30 : Inf.Coinf | 8.36E-01 | 1.41E-02 | 8.06E-01 | 8.62E-01 |
| Temp.HW : Inf.Coinf | 2.78E-01 | 1.75E-02 | 2.45E-01 | 3.13E-01 |

**Table S5. A model of LC host mortality indicated clear genotype and temperature effects, but their interaction did not drive mass mortality.** This model can be summarized as “Mortality ~ Temperature*Genotype + (1|batch)”. This model had a conditional Bayes R^2^ of 0.95 and a marginal of 0.94 (standard error of 0.002). We included a ten-level random effect of “Batch”. This model was for N2 and *srf-2* mutant *C. elegans*. Abbreviations are as described in Table S1, with “Geno.N2” for the N2 nematode genotype and “Geno.srf2” for the *srf-2* mutant genotype. Interaction terms are given by a colon. 95CIs that do not include zero are shown in bold. The coefficient values are logits.

| **Independent variable** | **Coefficient** | **Error** | **95CI, low** | **95CI, high** |
| --- | --- | --- | --- | --- |
| Intercept (Temp.20 : Geno.N2) | -5.19 | 0.34 | -5.90 | -4.54 |
| **Temp.25** | **0.85** | **0.38** | **0.13** | **1.62** |
| **Temp.30** | **4.13** | **0.32** | **3.54** | **4.80** |
| **Temp.HW** | **3.21** | **0.32** | **2.62** | **3.89** |
| **Geno.srf2** | **3.32** | **0.40** | **2.55** | **4.13** |
| Temp.25 : Geno.srf2 | -0.46 | 0.40 | -1.27 | 0.30 |
| Temp.30 : Geno.srf2 | -0.43 | 0.34 | -1.14 | 0.21 |
| **Temp.HW : Geno.srf2** | **-1.08** | **0.34** | **-1.78** | **-0.44** |
| Random effect (Batch) | 0.35 | 0.11 | 0.19 | 0.63 |

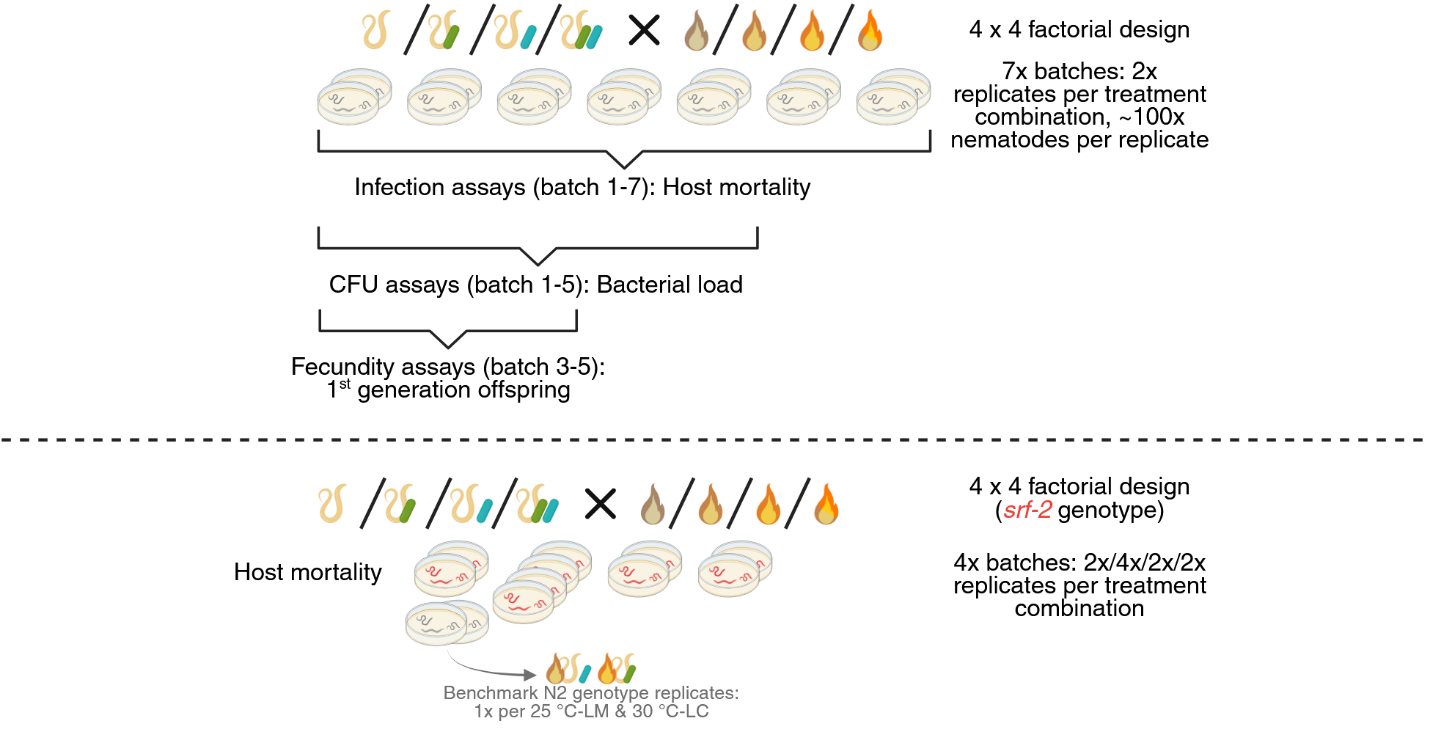

**Figure S1. Using a fully factorial design, we analyzed up to seven batches of N2 genotype infection assays to produce the primary dataset.** All replicates were assayed for mortality (n = 14). Nested CFU and fecundity assays indicate replicates were also used to collect bacterial load or offspring data, respectively. Separately, we collected host mortality data from *srf-2* mutant infection assays. In the first batch of *srf-2* experiments, we included single benchmarking replicates of N2 infection assays at peak virulence conditions (25 °C-LM and 30 °C-LC) and verified mortality was within the range of the primary N2 dataset. Figure created with Biorender.

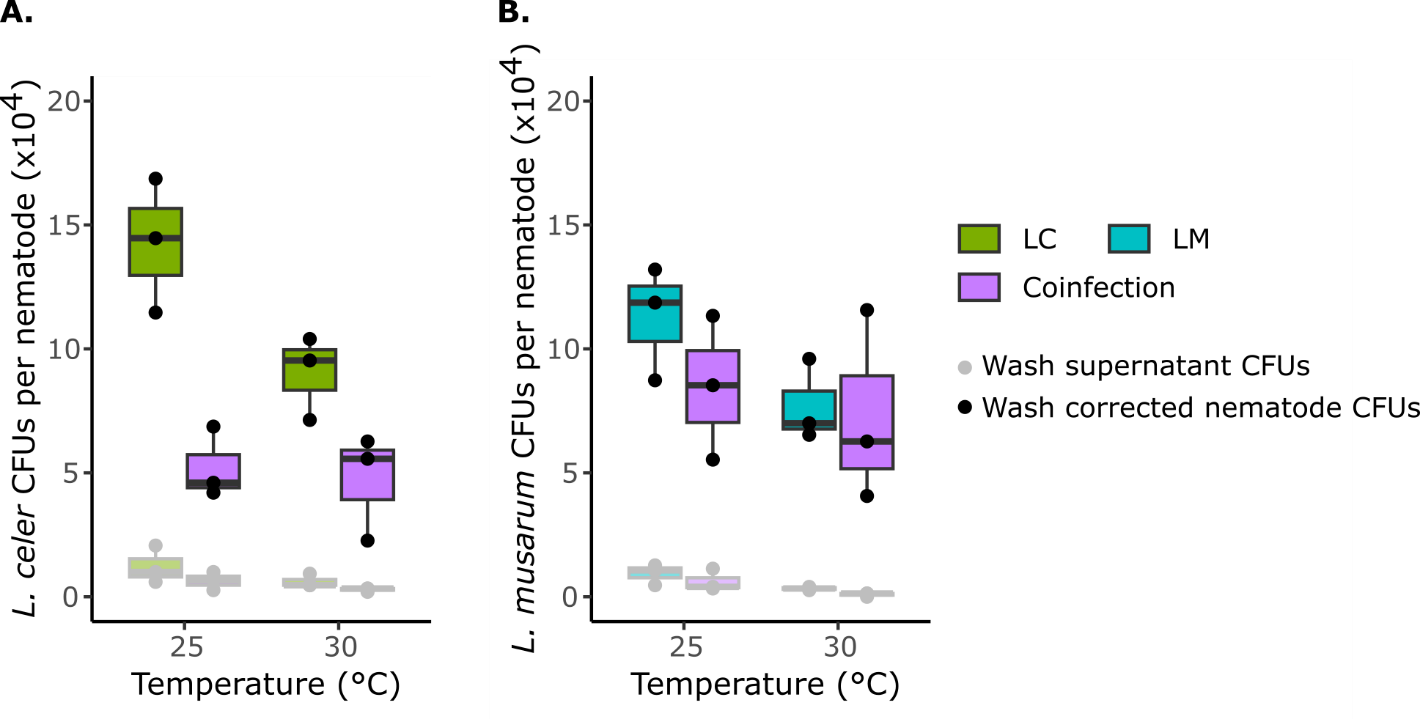

**Figure S2. Nematode washing removed the vast majority of bacteria not directly associated with the host.** We selected to benchmark the CFU assays only at 25 °C (peak LM CFUs, peak LM virulence) and 30 °C (minimum LC CFUs, peak LC virulence). For both *L. celer* (**A.**) and *L. musarum* (**B.**) washing appears effective (wash supernatant was ca. 10% of loads on nematodes) and therefore reported CFUs reflect trends of host-associated bacterial loads. The dark colored boxes (“Wash corrected”) CFUs are calculated by subtracting the “Wash supernatant” (light colored) values from the the total CFUs counted per replicate. We also measured host mortality for each replicate (not shown), validating that the infection assays were generally comprable to the data presented in the main text.

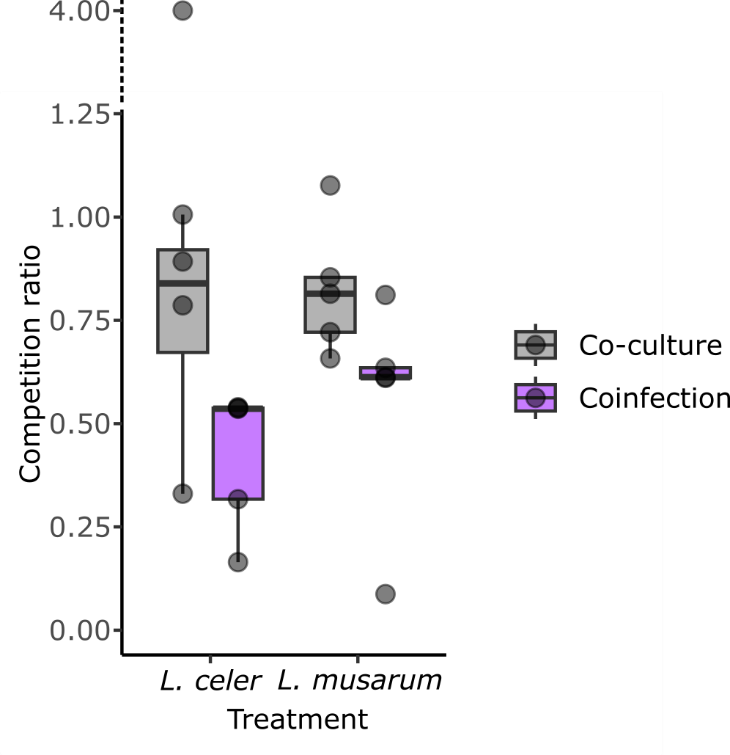

**Figure S3. Parasite coinfection competition mediated by the host appears to be stronger than environmental competition alone.** Co-culture (enviornmental competition alone, at 25 °C) made minor reductions in *L. celer* (11% median reduction) and *L. musaurm* (19% median reduction). While coinfection (on hosts at 25 °C) made greater reductions for both *L. celer* (46% median reduction) and *L. musaurm* (39% median reduction). “Competition ratio” is defined as the ratio of single-culture or infection CFUs to co-culture or coinfection CFUs per *Leucobacter* species, per plate (co-culture) or per nematode (co-infection). With caution for small samples sizes and a seemingly extreme value measured for *L. celer* CFUs on plates, we found statistical support for a difference of the effect of co-culture or coinfection treatments on *L. celer* CFU abundance (Kruskal-Wallis non-parametric test, chi-squared value = 3.94, p-value = 0.05). Data for *L. musarum* appeared to have a near normal distribution, and we therfore used a parametric Welch’s t-test, finding a nearly but not clearly statistical difference (t = 1.93, p-value = 0.10). Applying a Kruskal-Wallis test with *L. musarum* data to match the analysis conducted for *L. celer* yielded a “signficant” result (chi-squared value = 4.81, p-value = 0.03). Given the sample size of this supplemental analysis and data trends (median reductions), we inferred that parasite comeptition is likely excaerbated by interactions on the host. Co-culture plates were equivelent to infection assay plates immediately prior to the addition to nematodes (i.e., co-culture with OP50 on NGM agar for 18 h at 25 °C). Although infection assays also include an additional 30 h at varying temperatures with the addition of nematodes, we considered that we would not be able to properly disentangle the effects of nematode activity on those lawns (e.g., preferential eating of bacteria, shedding new bacteria on to the plate, etc.), such that these additional assays would not provide additonal insights. One batch of the infection assays used for mortality data was conducted in parallel with the first batch of these co-culture compeititon plate assays as a benchmark for experimental consistency. Note the broken y-axis to include a single high value point for *L. celer*.

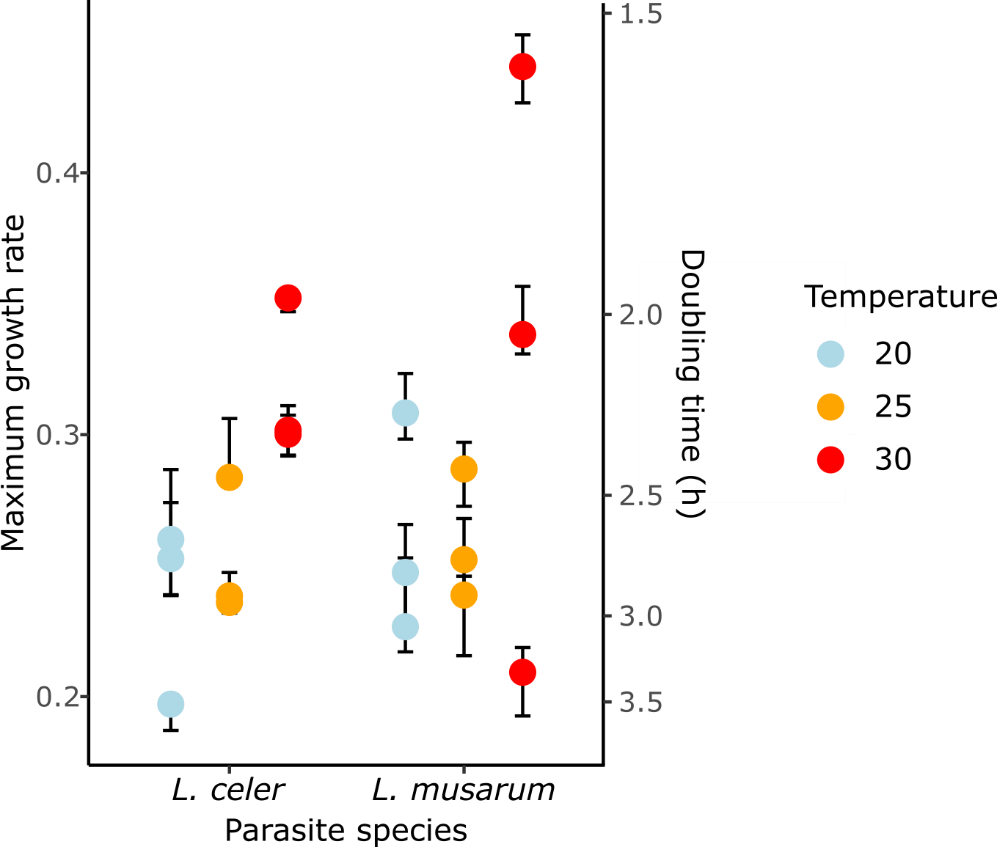

**Figure S4. Maximum growth rate observed in single cultures of *Leucobacter* grown *in vitro* at different temperatures.** Both genomic predictions and these experimental data suggest modestly increased maximum growth rates at approximately 30 °C compared to 20 or 25 °C. Points are median values of technical replicates (n =24) per batch (well plate) with 95% confidence interval error bars.

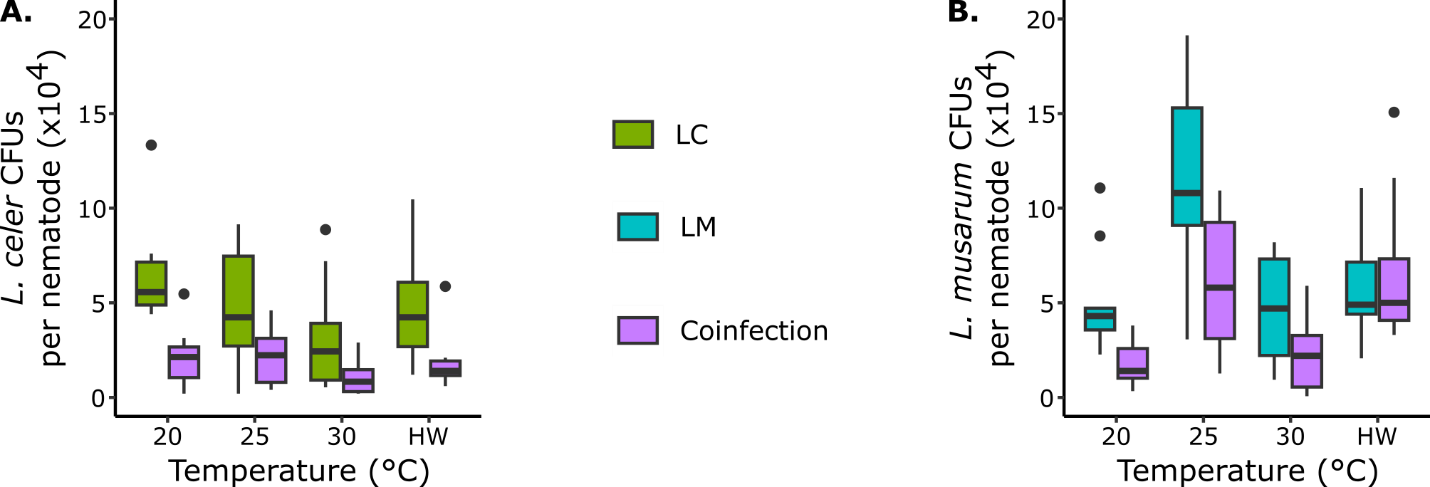

**Figure S5. Parasite fitness (CFU load) was generally reduced by coinfection, with secondary effects of temperature.** (**A.**) *Leucobacter celer* load was reduced by high temperature (30 °C) and by coinfecting *L. musarum*. (**B.**) *Leucobacter musarum* load peaked at 25 °C and was typically suppressed by coinfection. However, during heatwave, *L. musarum* loads were not reduced by coinfection. Heatwave is abbreviated as HW.

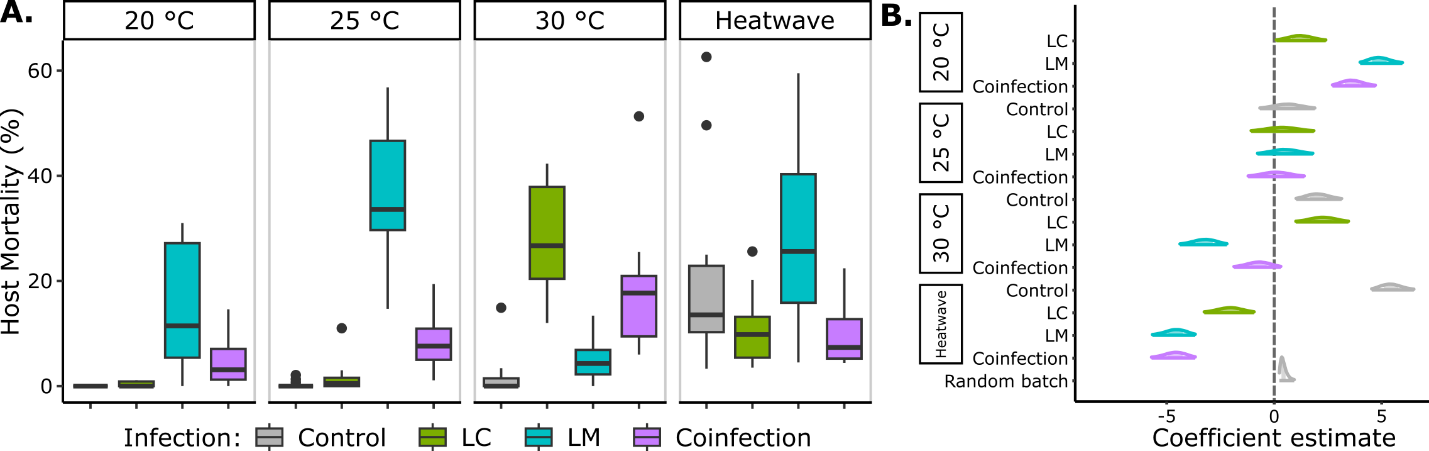

**Figure S6. Coinfecting parasite interactions modified temperature-dependent host mortality.** (**A.**) Boxplots of host mortality data for each infection treatment type faceted by temperature treatment. LM (*L. musarum* alone) and LC (*L. celer* alone) treatments with extreme warming (30 °C) reversed in relative severity. However, at all temperatures, coinfection protected hosts from peak mortality of the most harmful single infection. (**B.**) Posterior distributions of coefficient estimates from the GLMM, showing the 95CI. Negative model effects during 30 °C and heatwave indicated that mortality was partly due to temperature alone, with these infections being less severe than at cooler temperature. Control nematodes at 20 °C gave the intercept of this model, such that estimates of non-control infections at temperatures other than 20 °C represent interactive effects. Coefficients are on the logit scale.

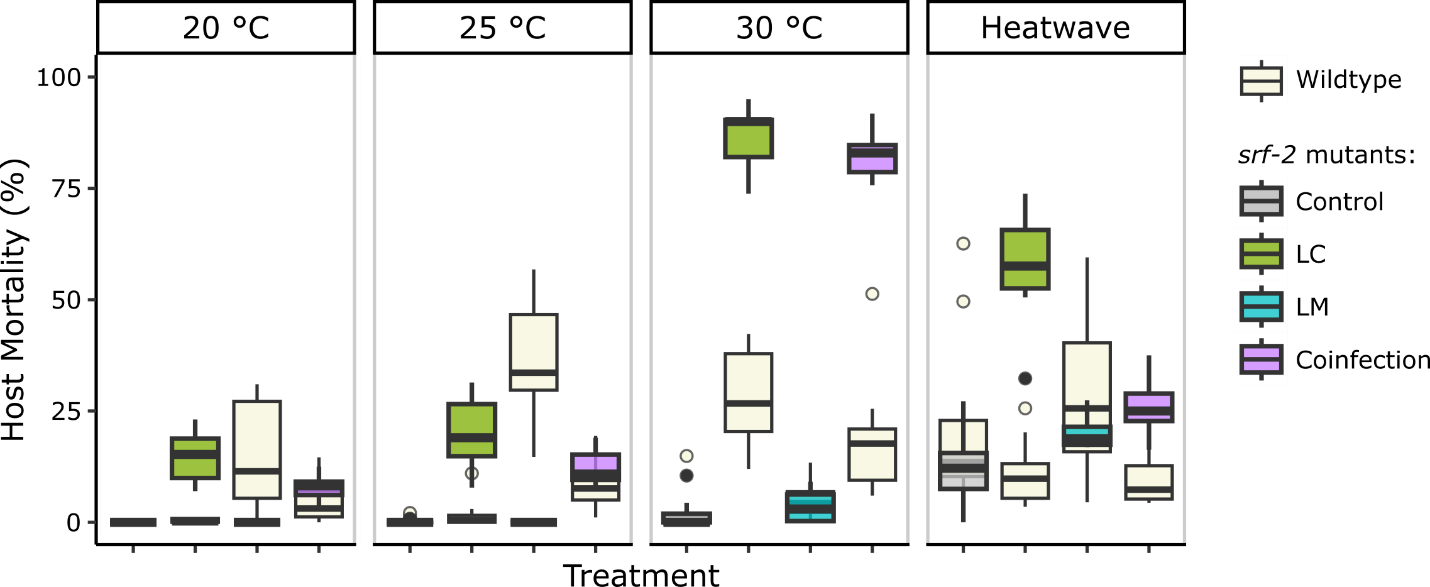

**Figure S7. Parasite interactions persisted to give intermediate mortality, despite major changes in host condition due to genetic susceptibility.** Colored boxes are infection treatment types on *srf-2* mutant hosts and the beige boxes are N2 wildtype data (as given in Figure S6). Mutant nematodes were more susceptible to *L. celer* infections across temperatures, with mass mortality observed at 30 °C. Parasite competition protected hosts across temperatures and with inversed host-parasite susceptibility. However, protection was notably less effective with combined thermal stress and genetic susceptibility (LC, *srf-2*, and 30 °C combination).

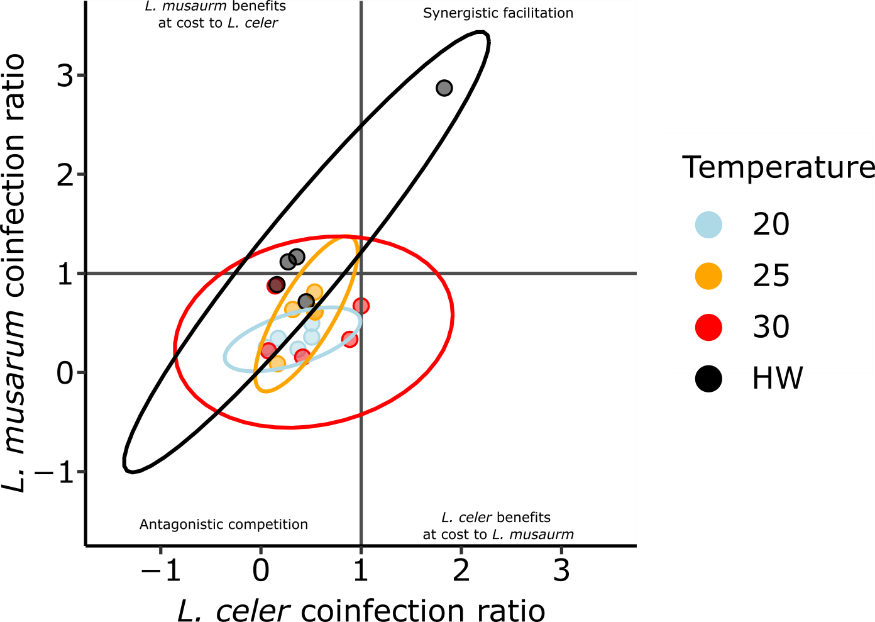

**Figure S8. Bacterial competition is typically mutually harmful.** The strength of coinfection competition per parasite species is given as the ratio of load during coinfection to load during single infection. Values greater than one indicate increased loads (positive effect of coinfection) and values less than one indicate reduced parasite loads (negative effect of coinfection). Load data per batch were averaged (two replicates), such that each data point represents one experimental batch. Coinfection ratios cannot be negative, but 95% confidence ellipses are given that span negative values.

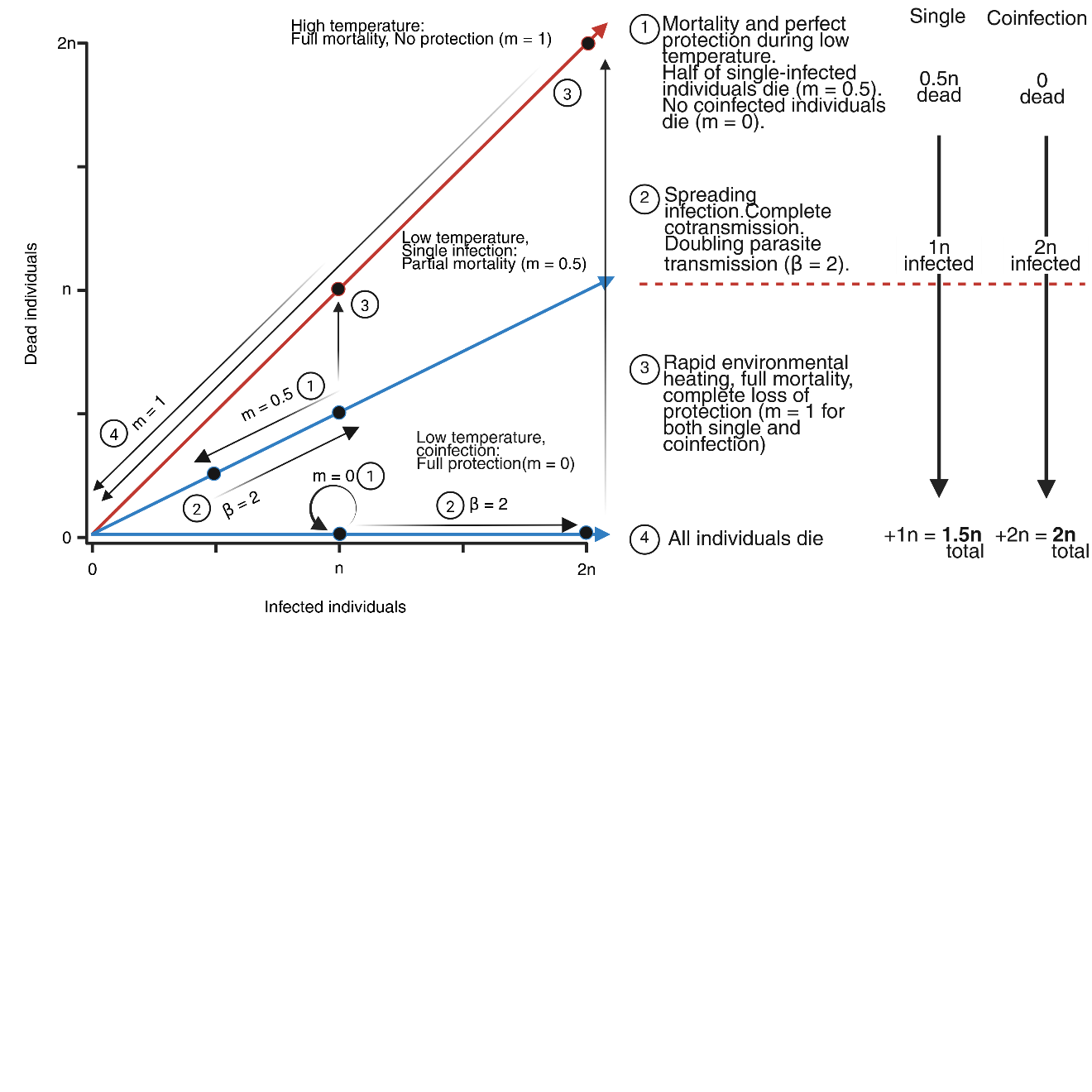

**Figure S9.** **Increased survivorship due to protection could lead to greater parasite transmission and downstream increases in deaths.** We briefly considered a hypothetical scenario where the temperature-dependent loss of protection during coinfection could lead to greater death throughout a population over time than consistent high levels of mortality. In this simple proof of concept, we defined mortality as the proportion of the infected population that dies as m (0 to 1). We set coinfection to completely protect (m = 0) and single infection as moderate (m = 0.5) at low temperature, with high temperature infections being fully lethal with total loss of protection (m = 1 for both single and coinfection). We additionally specified that infections spread in a temperature independent manner, such that transmission (β) is always greater than 1, which we set as β = 2 for simplicity. In such a scenario, the number of individuals that are infected reach the highest levels during low temperature coinfection, but then all die during heating – causing the greater total loss in the population. This outcome requires a sufficiently high level of protection during low temperature and transmission, paired with a significant breakdown of coinfection protection at high temperatures. For the experimental data which we collected during infection of *srf-2* hosts, 25 °C median mortality (e.g., low temperature m) was 0.19 for LC and 0.11 for coinfection. During 30 °C, m was 0.90 and 0.83, respectively. Based on these mortality rates, a transmission rate of β > 8.25 would be required to achieve net negative impacts of coinfection.

**Supporting document S1**

*Host fecundity is primarily shaped by temperature and secondarily by infection*

*Methods:* We assessed host fecundity as the number of hatched offspring produced by adults immediately following the infection assays. Using the washed nematodes collected from infection assays, we transferred three individuals in a M9 droplet with 200 µg/mL amoxicillin to a 6 cm NGM-OP50 food plate. This concentration of amoxicillin impairs *Leucobacter* growth without detrimental effects on the host (Clark & Hodgkin 2015; Yao *et al.* 2022). We pipetted these droplets at the edge of the plate, away from the bacterial lawn and observed that the nematodes could not immediately escape from the droplets. We incubated these plates at 20 °C. We counted all nematodes after 40 h of incubation. Our approach to measure the acute impact of infection differs from a previous measurement of host fecundity where hosts were exposed to *Leucobacter* throughout their adult lives (Hodgkin et al., 2013).

*Results:* We measured reductions of host fecundity as a sub-lethal fitness cost, in contrast to mortality. Fecundity could show the same trends as mortality or compensatory effects/terminal investment (Gulyas & Powell, 2022; Pike et al., 2019), possibly with temperatures-specific interactions not observed for mortality. To test if temperature and infection affected the reproductive potential of hosts that survive infection, we constructed a model of host fecundity (Supp. doc. S1 - Figure 1, Supp. doc. S1 - Table 1). This GLMM can be summarized as “Fecundity ~ Temperature*Infection + (1|Batch),” with a Nakagawa’s pseudo-R^2^ of 0.900 (conditional) and 0.895 (marginal). We observed a primary effect of temperature, with a secondary effect of infection (Supp. doc. S1 - Table 1).

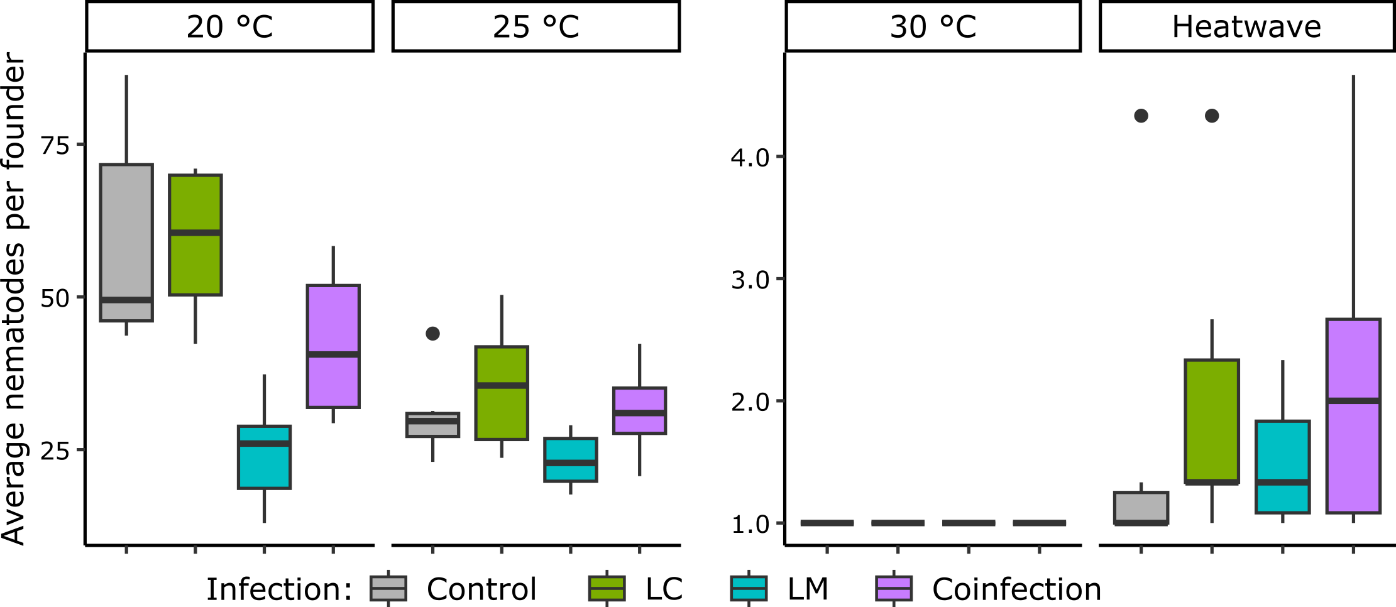

**Supp. doc. S1 - Figure 1. Temperature constrained host fecundity, with virulence impacts only when thermal stress was low.** Shown are the average number of nematodes on lineage assay plates after 40 h (first generation only) per founding nematode taken from infection assays. Combining moderate thermal and disease stress (25 °C LM) did not further worsen fecundity losses. At extreme levels of thermal stress, host reproduction essentially halted, precluding virulence effects. Note the difference in scale between 20-25 °C and 30 °C-heatwave plots.

**Supp. doc. S1 - Table 1. A model of host fecundity indicated clear temperature, infection, and interactive effects.** This frequentist GLMM (negative binomial) can be summarized as “Fecundity ~ Temperature *Infection + (1|Batch) + offset(Founders)”. In a single replicate, one of three nematodes died early in the analysis and we adjusted counts by the Founders offset term. The model had a Nakagawa’s pseudo-R^2^ of 0.900 (conditional) and 0.895 (marginal). Batch was a three-level random effect. Abbreviations are as described in Table S1. Effects with significant p-values (≤ 0.05) are in bold; non-significant effects at p ≤ 0.10 are underlined.

| **Independent variable** | **Coefficient** | **Error** | **Test statistic (z)** | **p-value** |
| --- | --- | --- | --- | --- |
| Intercept (Temp.20 : Inf.Con) | 2.17 | 0.12 | 18.74 | < 2e-16 |
| **Temp.25** | **-0.66** | **0.14** | **-4.82** | **1.42e-06** |
| **Temp.30** | **-4.08** | **0.27** | **-15.15** | **< 2e-16** |
| **Temp.HW** | **-3.61** | **0.23** | **-15.91** | **< 2e-16** |
| Inf.LC | 2.73e-04 | 0.13 | 0.002 | 0.998 |
| **Inf.LM** | **-0.87** | **0.14** | **-6.30** | **3.02e-10** |
| Inf.Coinf | -0.23 | 0.14 | -1.69 | 0.092 |
| Temp.25 : Inf.LC | 0.15 | 0.19 | 0.76 | 0.445 |
| Temp.30 : Inf.LC | -2.81e-04 | 0.38 | -0.001 | 0.999 |
| Temp.HW : Inf.LC | 0.22 | 0.31 | 0.71 | 0.479 |
| **Temp.25 : Inf.LM** | **0.61** | **0.20** | **3.08** | **0.002** |
| **Temp.30 : Inf.LM** | **0.87** | **0.38** | **2.28** | **0.024** |
| **Temp.HW : Inf.LM** | **0.80** | **0.33** | **2.46** | **0.014** |
| Temp.25 : Inf.Coinf | 0.25 | 0.19 | 1.30 | 0.193 |
| Temp.30 : Inf.Coinf | 0.23 | 0.38 | 0.60 | 0.549 |
| Temp.HW : Inf.Coinf | 0.55 | 0.31 | 1.79 | 0.073 |
| Random effect (Batch): 0.014 variance, 0.117 standard deviation | | | | |

Temperature primarily constrained host fitness potential, but infection reduced fecundity in the absence of thermal stress. Control nematodes produced the most offspring near their reproductive optimum at 20 °C, with reduced fecundity at 25 °C, no offspring at 30 °C (as observed in Begasse et al. 2015), and only few offspring following the heatwave exposure (Supp. doc. S1 - Figure 1, Supp. doc. S1 - Table 1). Host fecundity across infections trended as host mortality when at the host thermal optimum (20 °C) (Figure 2). That is, LM significantly reduced host fitness (effect = -0.87, z = -6.30, p < 0.001), LC treatments had no meaningful effect, and coinfection had an unclear intermediate negative effect (effect = -0.23, z = -1.68, p = 0.092).

Additional stressors did not necessarily drive additional fecundity reductions. Control nematodes at 25 °C (i.e., temperature stress alone) and LM infected hosts at 20 °C (i.e., infection stress alone) both had reduced fitness compared to control nematodes at 20 °C (Supp. doc. S1 - Figure 1, Supp. doc. S1 - Table 1). However, the combination of both LM and 25 °C did not further reduce fecundity (with a “balancing” positive interactive effect = 0.61, z = 3.08, p = 0.002). Extreme thermal stress precluded any potential negative impacts of infection. With no/few offspring produced at 30 °C and during heatwave, negative LM treatment effects on fecundity were prevented (Supp. doc. S1 - Table 1). Coinfected nematodes may have produced slightly more offspring than controls during heatwave (effect = 0.55, z = 1.79, p = 0.073) (Supp. doc. S1 - Figure 1, Supp. doc. S1 - Table 1). This trend is in line with our observations that coinfection and LC treatment possibly reduced the effects of thermal stress on host mortality (Figure 2, Table S1).

*Discussion:* These results suggest that temperature and infection have the potential to mask, rather than amplify, each other. Reduced host fecundity at 25 °C corresponds to a thermal tipping point that may be linked to lower rates of fertilization (Begasse et al., 2015; Gouvêa et al., 2015; Harvey & Viney, 2007; Petrella, 2014). If virulence also lowers fertilization rates, we reason that temperature and disease cannot combine to reduce fecundity below a bottom limit of halting fertilization (but importantly not killing already formed embryos). At higher experimental temperatures, cessation of host reproductive capacity was essentially complete, precluding meaningful additional impacts of infection. Physiological limitations (i.e., no less than zero fertilizations or viable offspring possible) may have thus masked the dual impacts of heating and disease on host reproduction. Hosts may generally benefit from acquiring parasites providing protection at less stressful temperatures (e.g., *L. celer*), when high mortality from the protector only occurs at severe temperatures that end host fitness regardless.

Previous work has demonstrated shifts in fitness tradeoffs at the gene expression level between immunity and reproduction in *C. elegans* infected or protected by *Leucobacter* (Will et al., 2025). From those transcriptomic data (at 25 °C), we inferred the costliest immunity-reproduction tradeoff to occur during LM infection, which are reduced during protective coinfection. Here, we correlated mortality and reproduction to show that there appears to be tradeoffs at the phenotypic level as well (Supp. doc. S1 - Figure 2)

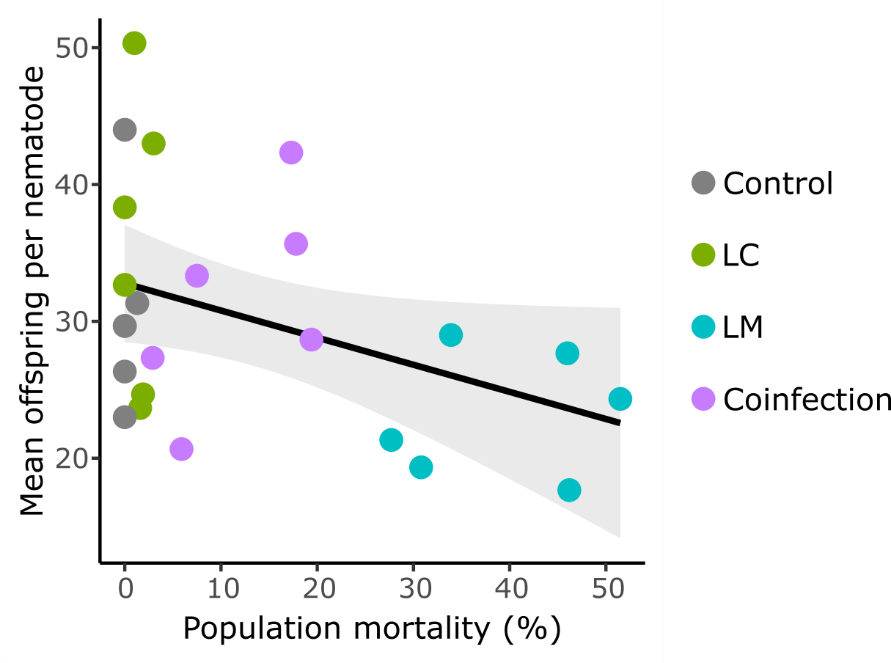

**Supp. doc. S1 - Figure 2. Reported immune-reproduction transcriptomic tradeoffs appear to correspond to phenotypic effects.** At 25 °C, we found fecundity was most reduced in nematodes during LM treatment. LM-25 °C also elicits the strongest upregulation of immune genes with strongest downregulation of reproduction genes, as found in (Will et al., 2025). Overall, these data indicate a negative relationship between disease severity and reproductive potential (Pearson’s correlation = -0.4, p = 0.05), to which we infer a combination of high immune investment and morbidity contribute. Although these data sets (RNAseq or phenotypes) come from parallel experiments, the trend suggested that gene expression changes reflect biologically relevant changes in reproduction.

**Supporting document S2**

*Model*

Here, we describe the mathematical model in more detail. As a reminder, we focus on a population which is partitioned into four distinct groups depending on their disease status. There are two infecting parasites, which we denote as Parasite 1 (P1, *Leucobacter celer*) and Parasite 2 (P2, *Leucobacter musarum*), and when both are present in the environment, hosts may be coinfected. We denote our populations at some time $t$ as $S(t)$ for the susceptible population, $I_{1}(t)$ and $I_{2}(t)$ for the populations infected with P1 and P2 respectively, and $C\left( t \right)$ the coinfected population. We make several simplifying assumptions to ensure that we can attribute any observed behaviour to the key processes. Firstly, we will assume that the average reproductive period is much longer than the infectious period so that we can neglect any births. Further, we assume that all hosts undergo a small amount of natural, non-disease-related mortality, which is the same for all host types. We will model a saturated environment, meaning that the parasites will be in abundance so that we do not need to track their densities in the environment. As such, transmission rates $\beta_{1}$ and $\beta_{2}$ for P1 and P2 respectively are interpreted as the rates of establishing an infection of the associated type. Each parasite is able to induce mortality related to infection, with rates $\alpha_{1}$ and $\alpha_{2}$ (virulence).

Finally, when modelling the simultaneous presence of both parasites, they may interact in a way which benefits the host. There are two mechanisms which can do this: exclusion and attenuation. Exclusion is defined to be the reduction in infectiousness of one parasite when already infected with the other. We define the strength of exclusion conferred to a host with P1 against P2 as $y_{e1}\in[0, 1]$, where 0 is no exclusion and 1 is complete exclusion. This acts on the transmission of P2 to hosts with P1 through a decreasing function $E_{1}:\left[ 0,1 \right]\to[0,1]$, so that $E_{1}(y_{e1})$ is the total loss of transmission caused by P1 against P2 with a strength $y_{e1}$. There are similar definitions for exclusion conferred to hosts with P2 against P1 ($y_{e2}$ and $E_{2}$). For simplicity, we will assume that:

$$E_{i}\left( y_{ei} \right)=1-y_{ei} \mathrm{for}i=1,2.$$

(S1)

Attenuation only acts on hosts which are coinfected, and protects against the adverse effects of having both parasites. In this case, we will write the co-virulence term as a function $A:\left[ 0, 1 \right]^{2}\to\mathbb{R}_{+}$, which takes the form

$$A\left( y_{a1},y_{a2} \right)=\left( 1-y_{a2} \right)\alpha_{1}+\left( 1-y_{a1} \right)\alpha_{2},$$

(S2)

where $y_{a1}$ is the level of attenuation conferred to the host by P1 against P2, with a similar definition for $y_{a2}$. Note that we assume that under no attenuation, the effects of having both parasites is additive for virulence.

These assumptions combine to yield the following system of ordinary differential equations (ODEs):

| $\frac{dS}{dt}=-\left( d+\beta_{1}+\beta_{2} \right)S,$ | $S\left( 0 \right)=N_{0,}$ | (S3) |
| --- | --- | --- |
| $\frac{dI_{1}}{dt}=\beta_{1}S-\left( d+\alpha_{1}+E_{1}\left( y_{e1} \right)\beta_{2} \right)I_{1},$ | $I_{1}\left( 0 \right)=0,$ | (S4) |
| $\frac{dI_{2}}{dt}=\beta_{2}S-\left( d+\alpha_{2}+E_{2}(y_{e2})\beta_{1} \right)I_{2},$ | $I_{2}\left( 0 \right)=0,$ | (S5) |
| $\frac{dC}{dt}=E_{1}(y_{e1})\beta_{2}I_{1}+E_{2}(y_{e2})\beta_{1}I_{2}-\left( d+A(y_{a1}, y_{t2}) \right)C.$ | $C\left( 0 \right)=0.$ | (S6) |

*Mortality*

We will define mortality $M(t)$ as the total proportion of hosts that die by time $t$ of the simulation. Since we have no births into the system, the density of hosts that die is precisely the initial population size minus those still alive, so that if we write the total population size to be $N\left( t \right)=S\left( t \right)+I_{1}\left( t \right)+I_{2}\left( t \right)+C(t)$:

$$M\left( t \right)=\frac{N_{0}-N\left( t \right)}{N_{0}}=1-\frac{N\left( t \right)}{N_{0}}.$$

(S7)

We differentiate both sides of this to obtain an ODE for the mortality in time. The value of the derivative of the total population size can be found by adding together the four ODEs (S3)-(S6), to yield the following expression for the derivative of the mortality:

| $\frac{dM}{dt}=\frac{dS+\left( d+\alpha_{1} \right)I_{1}+\left( d+\alpha_{2} \right)I_{2}+\left( d+\left( 1-y_{a2} \right)\alpha_{1}+\left( 1-y_{a1} \right)\alpha_{2} \right)C}{N_{0}},$ | $M\left( 0 \right)=0.$ | (S8) |
| --- | --- | --- |

*Single Infection*

To evaluate the effects of temperature when only a single parasite is present, we will write a truncated version of the model which considers only a single infective class (and hence no coinfected class). Here, we will consider P1 being present and P2 not, but similar expressions exist for P2. Firstly, we now have a truncated ODE system:

| $\frac{dS}{dt}=-\left( d+\beta_{1} \right)S,$ | $S\left( 0 \right)=N_{0},$ | (S9) |
| --- | --- | --- |
| $\frac{dI_{1}}{dt}=\beta_{1}S-\left( d+\alpha_{1} \right)I_{1},$ | $I_{1}\left( 0 \right)=0.$ | (S10) |

We are able to solve this system analytically for all times, with the following expressions denoting the solution:

$$S\left( t \right)=N_{0}e^{-\left( d+\beta_{1} \right)t},$$

(S11)

$$I_{1}\left( t \right)=\left\{ \begin{aligned} \frac{\beta_{1}N_{0}}{\beta_{1}-\alpha_{1}}\left[ e^{-\left( d+\alpha_{1} \right)t}-e^{-\left( d+\beta_{1} \right)t} \right], &\beta_{1}\neq\alpha_{1}, \\ \beta_{1}N_{0}te^{-\left( d+\beta_{1} \right)t}, &\beta_{1}=\alpha_{1}\neq0, \end{aligned} \right.$$

(S12)

which yields the following expression for the mortality proportion:

$$M\left( t \right)=1-\frac{N\left( t \right)}{N_{0}}=\left\{ \begin{aligned} 1-\frac{\beta_{1}}{\beta_{1}-\alpha_{1}}e^{-\left( d+\alpha_{1} \right)t}+\frac{\alpha_{1}}{\beta_{1}+\alpha_{1}}e^{-\left( d+\beta_{1} \right)t}, &\beta_{1}\neq\alpha_{1}, \\ 1-\left( 1+\beta_{1}t \right)e^{-\left( d+\beta_{1} \right)t}, &\beta_{1}=\alpha_{1}. \end{aligned} \right.$$

(S13)

It can be seen that if we consider the mortality (S13) as a function of $\beta_{1}$ and $\alpha_{1}$, then the contours $M\left( t \right)=m$ in $\left( \beta_{1},\alpha_{1} \right)$-space are monotonically decreasing, convex functions with the following asymptotic properties (see Figure 1):

$$\alpha_{1}\to d-\frac{1}{t}\ln\left( 1-m \right) \mathrm{as} \beta_{1}\to\infty,$$

(S14)

$$\beta_{1}\to d-\frac{1}{t}\ln\left( 1-m \right) \mathrm{as} \alpha_{1}\to\infty.$$

(S15)

*Simulation*

In this section, we describe the simulation methods, and in particular, the methods for the coinfection results. Firstly, we fix the final time to be $T$ and determine the mortalities from the single infections data, and set them according to the experimental data as follows: at ${20}^{\circ}C$, the mortality of P1 is $m_{1}=0.004$ and the mortality for P2 is $m_{2}=0.148$, for ${25}^{\circ}C$ we use $m_{1}=0.151$ and $m_{2}=0.369$, and for ${30}^{\circ}C$ we use $m_{1}=0.282$ and $m_{2}=0.048$. For each temperature, we find $(\beta_{j},\alpha_{j})$ pairs for each microbe using the mortality equation (S13) by sampling 1000 points randomly across the contour, with $d=0.0001$ and $t=20$ fixed.

For each parameter combination $(\beta_{1},\alpha_{1},\beta_{2}\alpha_{2})$, we then sample 20 protection values $\left( y_{e1},y_{e2} \right)$ or $\left( y_{a1}, y_{a2} \right)$ uniformly at random from the unit square, and calculate the co-mortality, $M(t)$, using a numerical solution of equation (S7). This yields a dataset which is 20000 in size, which we then filter by ensuring that the value $M(t)$ lies between $p_{1}$ and $p_{2}$, which is defined as co-mortality. These results are then plotted. In Figure 6, each dot represents one of the parameter combinations $(\beta_{1}, \alpha_{1})$ for panel A and C, and $(\beta_{2}, \alpha_{2}$) for panel B and D, where at least one of the randomized protection pairs yields intermediate co-mortality. We summarized all of the protection pairs which have intermediate co-mortality using a kernel density estimate plot (simulated using kdeplot from the seaborn package in Python) (Supp. doc. S3 - Figure 1).

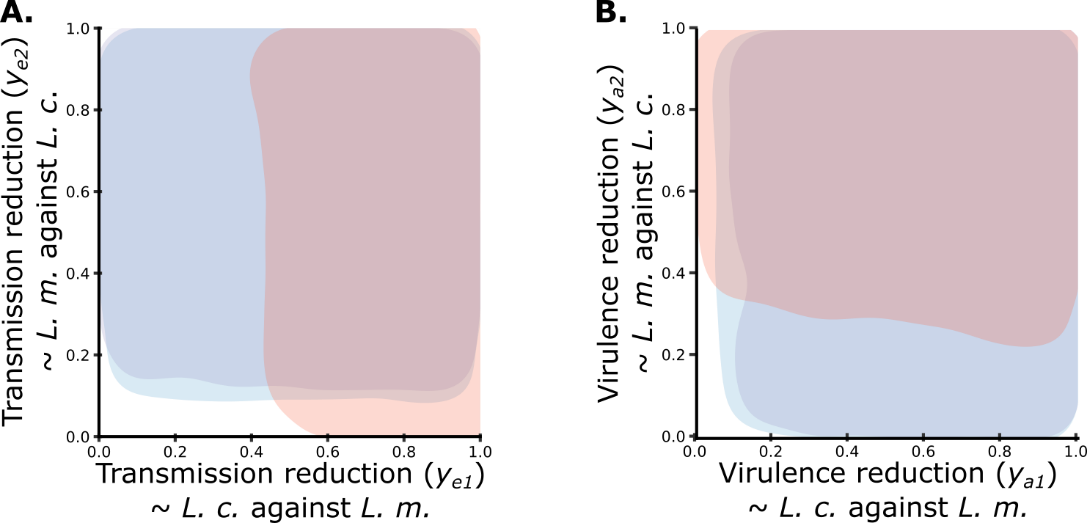

**Supp. doc. S3 - Figure 1. Each parasite could confer a range of protection levels (i.e., strength of competition), which was most limited by high thermal stress.** The shaded region for each colour is the region which contains 90% of the pairs which yield intermediate co-mortality for the various temperature regimes. The strength of competition (i.e., strength of protection) whether by exclusion (**A.**) or attenuation (**B.**) necessary to maintain intermediate mortality across temperature scenarios (overlap of blue, purple, and red zones) includes a broad range of values. High thermal stress (30 °C) is the most restrictive in both cases, corresponding experimentally to P2 *L. musarum* (*L. m.*) mediated protection against P1 *L. celer* (*L. c.*).

**Supporting document S3**

*Severe warming may link coinfecting parasite loads with virulence and protection*

Parasite loads and virulence were, tentatively, linked in a temperature by species manner. Having observed reduced parasite fitness (loads) and infection virulence during coinfection (see main text, Supp. doc. S3 - Figure 1), we tested if parasite loads could directly predict host outcomes. We constructed a Bayesian logistic regression that modeled host mortality during coinfection as a response to bacterial abundance and temperature (Supp. doc. S3 - Table 1). Implicit in this model is an assumption that parasite burden may drive host outcomes, rather than moribund hosts becoming differentially colonized. Connecting host and parasite fitness is inherently biologically relevant, therefore, we continued with this model despite inconclusive support using model selection methods (e.g., leave one out information criterion, LOOIC) (Supp. doc. S3 - Table 2). We found temperature-specific effects of parasite species loads acting in opposition to each other (Supp. doc. S3 - Table 1).

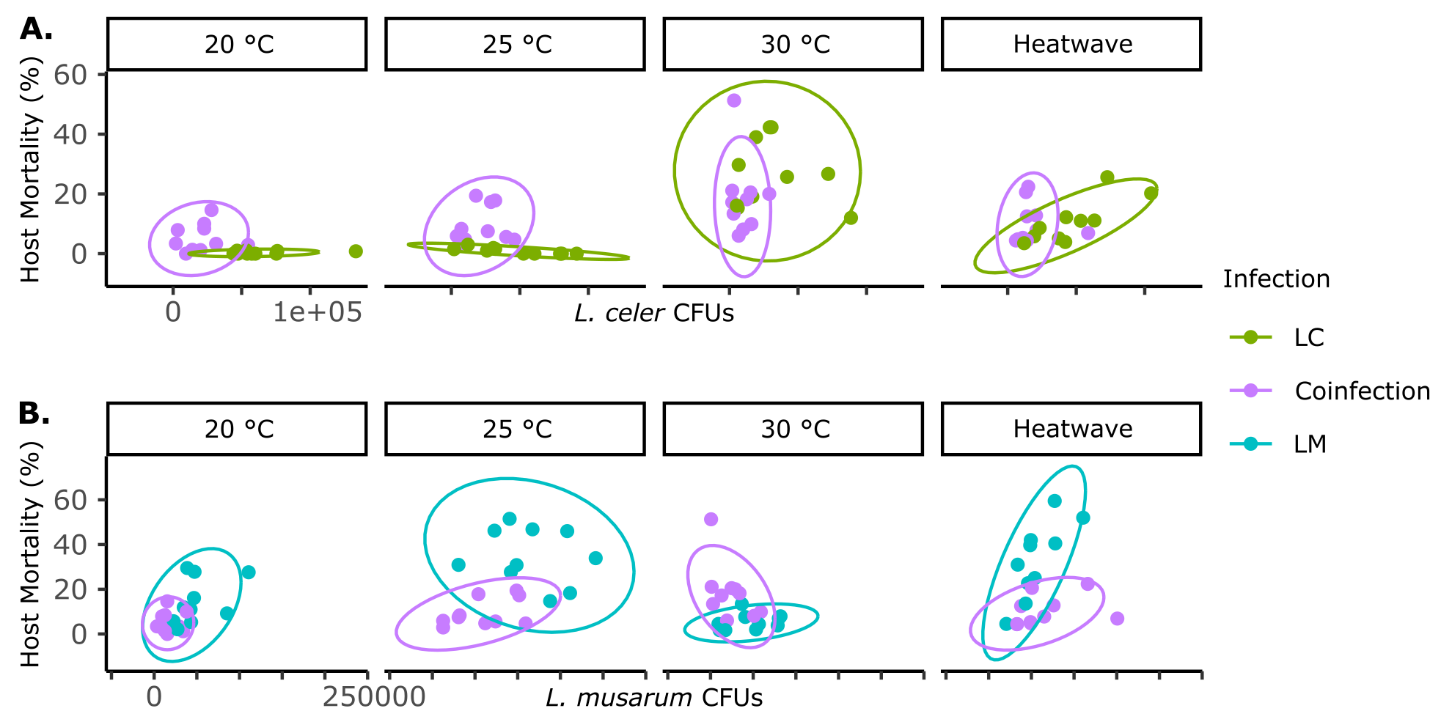

**Supp. doc. S3 - Figure 1. Parasite load and host mortality during coinfection**. For both *L. celer* (**A.**) and *L. musarum* (**B.**) parasite load and host mortality relationships had the most variation at peak virulence (i.e., 30 °C for LC and 25 °C for LM treatments). Ellipses are 95% confidence intervals.

**Supp. doc. S3 - Table 1. A model of coinfected host mortality indicated clear temperature-dependent effects of parasite load on infection virulence.** This Bayesian logistic regression can be summarized as “Mortality ~ CFU_*L.c.**Temperature + CFU_*L.m.**Temperature + (1|Batch)”, with CFU_*L.c.* and CFU_*L.m.* giving the CFU counts of *L. celer* and *L. musarum*, respectively*.* We centered scaled the bacteria CFU abundance data (raw *L. musarum* CFU counts were often approximately two-fold higher than *L. celer*). The model had a conditional (including random effects) Bayes R^2^ of 0.64 and a marginal (fixed effects only) of 0.43 (standard error of 0.03). Batch was a five-level random effect. This model was only for coinfection. Abbreviations are as described in Table S1. 95CIs that do not include zero are shown in bold. Less clear, weaker effects that exclude zero in the 90CI are underlined. The coefficient values are logits.

| **Independent variable** | **Coefficient** | **Error** | **95CI, low** | **95CI, high** |
| --- | --- | --- | --- | --- |
| Intercept (Temp.20) | -2.67 | 0.42 | -3.49 | -1.87 |
| CFU_*L.c.* | -0.11 | 0.15 | -0.42 | 0.18 |
| CFU_*L.m.* | 0.12 | 0.38 | -0.62 | 0.86 |
| Temp.25 | 0.24 | 0.30 | -0.35 | 0.85 |
| Temp.30 | 0.55 | 0.30 | -0.02 | 1.14 |
| Temp.HW | 0.24 | 0.32 | -0.38 | 0.88 |
| CFU_*L.c.* : Temp.25 | 0.12 | 0.20 | -0.26 | 0.52 |
| **CFU_*L.c.* : Temp.30** | **0.93** | **0.28** | **0.39** | **1.47** |
| CFU_*L.c.* : Temp.HW | -0.09 | 0.24 | -0.55 | 0.37 |
| CFU_*L.m.* : Temp.25 | 0.13 | 0.39 | -0.62 | 0.90 |
| **CFU_*L.m.* : Temp.30** | **-1.84** | **0.45** | **-2.71** | **-0.96** |
| CFU_*L.m.* : Temp.HW | 0.20 | 0.39 | -0.55 | 0.96 |
| Random effect (batch) | 0.63 | 0.37 | 0.24 | 1.59 |

**Supp. doc. S3 - Table 2. Leave one out information criterion (LOOIC) for models of host mortality using temperature and parasite load predictors.** The GLMMs of host mortality that included parasite load data as predictors was ranked marginally better or equivalent to than models that only included temperature predictors. The same order ranking was obtained if we used the Widely Applicable Information Criterion (WAIC, not shown). LOOIC model selection indicates that including parasite load data do not clearly construct more efficient models, with statistically-unclear improvements in LOOIC and expected log predictive density (ELPD), but appreciable improvement of Bayes R^2^. The coinfection models including temperature or load were not greatly improved over a null model – which highlights the stabilizing effect of coinfection protection (intermediate mortality) across temperatures. However, investigating a possible parasite load-infection virulence link is inherently biologically relevant, and therefore we present these analyses here. Both full single-infection models were of the basic form “Mortality ~ CFU*Temperature + (1|Batch)”.

| **Model** | **ELPD difference** | **ELPD error** | **LOOIC** | **R2 (cond.)** | **R2 (marg.)** |
| --- | --- | --- | --- | --- | --- |
| Coinfection, with species-specific loads | NA | NA | 371.9 | 0.644 | 0.427 |
| Without load | -13.2 | 25.2 | 398.3 | 0.465 | 0.271 |
| Null (~1) | -28.0 | 42.2 | 428.0 | 0.401 | 0.168 |
| LC, with load | NA | NA | 250.2 | 0.830 | 0.819 |
| Without load | -17.3 | 8.7 | 284.8 | 0.752 | 0.730 |
| Null (~1) | -321.7 | 51.8 | 893.6 | 0.096 | 0.070 |
| LM, with load | NA | NA | 412.0 | 0.865 | 0.687 |
| Without load | 0.0 | 18.1 | 412.1 | 0.835 | 0.529 |
| Null (~1) | -221.0 | 54.0 | 854.1 | 0.472 | 0.131 |

*Leucobacter celer* and *L. musarum* loads had contrasting effects on host mortality during coinfection at high temperature. Although coinfection lowered overall loads at every temperature, and consistently led to intermediate virulence, we only detected a clear per-CFU relationship between load and mortality with high thermal stress (30 °C). Indeed, during coinfection, models incorporating load and/or temperature did not strongly outperform a null model – in agreement with the stabilizing intermediate-mortality effect of protective coinfection we observed across temperatures. During coinfection at moderate temperatures or heatwave, incremental changes in parasite loads did not predict host mortality. However, with constant high temperatures (30 °C), we found patterns revealing load-dependent *L. celer* infection virulence in contrast to *L. musarum* load-dependent protection. We found coinfected *C. elegans* mortality increased with *L. celer* loads during 30 °C treatments (effect = 0.93, 95CI = 0.39, 1.47) (Supp. doc. S3 - Figure 2). In contrast, *L. musarum* abundance protected hosts against *L. celer* with a negative effect on mortality at 30 °C (effect = -1.84, 95CI = -2.71, -0.96) (Supp. doc. S3 - Figure 2). As *L. musarum* was generally more abundant than *L. celer* (main text results, Supp. doc. S3 - Figure 2), we attribute the finding of higher combined-parasite total loads predicating reduced mortality (see main text) to the preponderance of *L. musarum*.

Although model predictions for *L. musarum* loads during coinfection at non-30 °C temperatures appear to have a positive relationship (slope) with mortality (Supp. doc. S3 - Figure 2) and coefficient estimates were positive (Supp. doc. S3 - Table 1), the error surrounding these estimates was too great to conclude a load-mortality effect. Overall, these data may hint at broad load-mortality effects, but the statistics suggest a non-zero effect only at 30 °C.

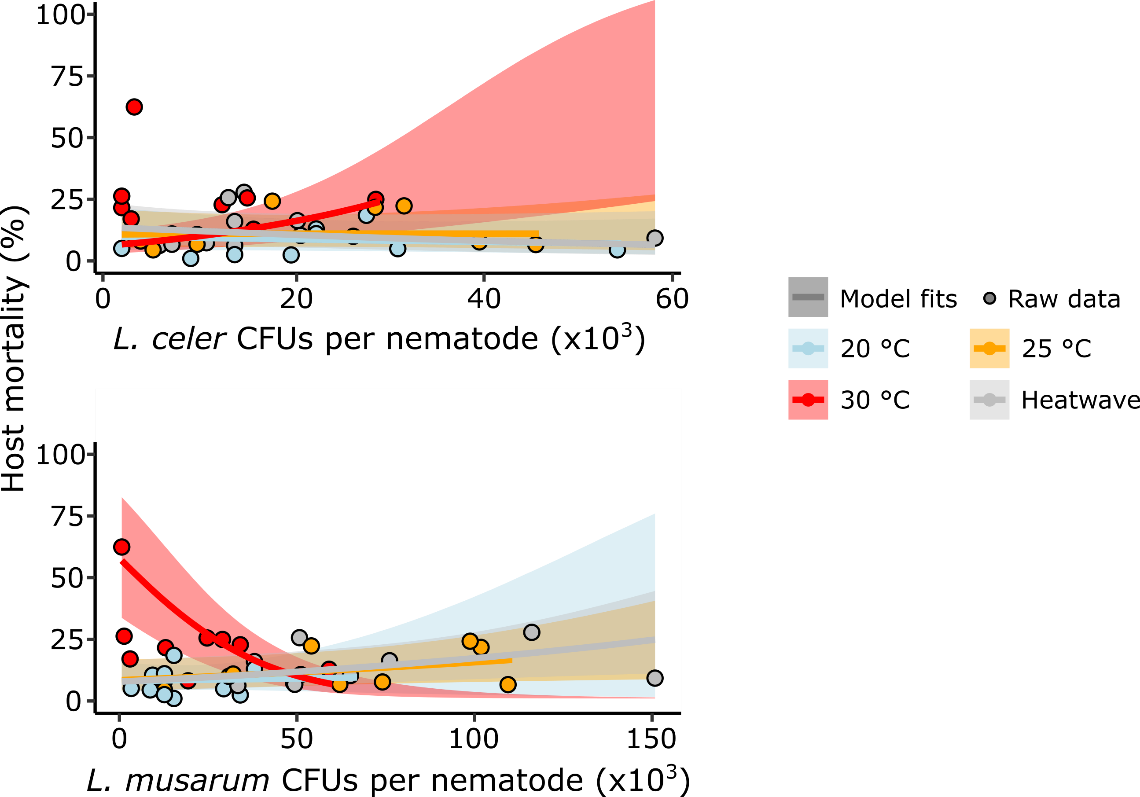

**Supp. doc. S3 - Figure 2. Bacterial loads during coinfection clearly relate to host mortality at 30 °C, with antagonistic effects of *L. celer* and *L. musarum*. A.** Model fit predictions of mortality during coinfection (lines with 95% confidence interval shading) were made across observed CFU abundance for the focal bacteria (x axis), with the coinfecting bacteria CFUs held at their mean value. The raw observed data have been plotted as points. We only observed clear effects of coinfecting parasite loads on virulence at 30 °C (red). Model estimate lines have been truncated at the maximum observed CFU value per treatment.

*Single infection load and mortality*

LC treatments showed no increases in mortality due to higher *L. celer* loads (Supp. doc. S3 - Table 3). LM treatments showed a modest load-mortality link that was maintained across temperatures, increasing in severity with heatwave (Supp. doc. S3 - Table 4). See Figure 2B in the main text for plotted raw data and model predictions. Although the model of LM host mortality suggested a load-mortality link (Supp. doc. S3 - Table 4), the inclusion of load did not notably improve model performance by LOOIC selection (Supp. doc. S3 - Table 2).

**Supp. doc. S3 - Table 3. A model of LC treatment host mortality did not find increasing parasite loads to increase mortality.** This Bayesian logistic regression can be summarized as “Mortality ~ CFU_*L.c.**Temperature + (1|Batch)”, with CFU_*L.c.* giving the CFU counts of *L. celer.* We centered scaled the bacteria CFU abundance data to be consistent with use of scaling in the coinfection model. The model had a conditional (including random effects) Bayes R^2^ of 0.83 and a marginal (fixed effects only) of 0.82. Batch was a five-level random effect. Abbreviations are as described in Table S1. 95CIs that do not include zero are shown in bold. Less clear, weaker effects that exclude zero in the 90CI are underlined. The coefficient values are logits.

| **Independent variable** | **Coefficient** | **Error** | **95CI, low** | **95CI, high** |
| --- | --- | --- | --- | --- |
| Intercept (Temp.20) | -5.15 | 0.47 | -6.12 | -4.30 |
| CFU_*L.c.* | -0.05 | 0.44 | -1.01 | 0.70 |
| Temp.25 | -0.43 | 0.70 | -1.86 | 0.88 |
| **Temp.30** | **3.99** | **0.45** | **3.17** | **4.94** |
| **Temp.HW** | **2.99** | **0.45** | **2.18** | **3.94** |
| **CFU_*L.c.* : Temp.25** | **-1.51** | **0.66** | **-2.82** | **-0.21** |
| CFU_*L.c.* : Temp.30 | -0.03 | 0.44 | -0.78 | 0.94 |
| CFU_*L.c.* : Temp.HW | 0.64 | 0.46 | -0.15 | 1.64 |
| Random effect (batch) | 0.28 | 0.22 | 0.03 | 0.84 |

**Supp. doc. S3 - Table 4. A model of LM treatment host mortality suggested possible load-mortality relationships, increasing in severity with heatwaves.** This Bayesian logistic regression can be summarized as “Mortality ~ CFU_*L.m.**Temperature + (1|Batch)”, with CFU_*L.m.* giving the CFU counts of *L. musarum*. We centered scaled the bacteria CFU abundance data to be consistent with use of scaling in the coinfection model. The model had a conditional (including random effects) Bayes R^2^ of 0.87 and a marginal (fixed effects only) of 0.69. Batch was a five-level random effect. Abbreviations are as described in Table S1. 95CIs that do not include zero are shown in bold. Less clear, weaker effects that exclude zero in the 90CI are underlined. The coefficient values are logits.

| **Independent variable** | **Coefficient estimate** | **Error** | **95CI, low** | **95CI, high** |
| --- | --- | --- | --- | --- |
| Intercept (Temp.20) | -1.66 | 0.38 | -2.43 | -0.91 |
| **CFU_*L.m.*** | **0.24** | **0.12** | **0.01** | **0.47** |
| **Temp.25** | **0.76** | **0.14** | **0.49** | **1.02** |
| **Temp.30** | **-1.16** | **0.18** | **-1.52** | **-0.82** |
| **Temp.HW** | **1.06** | **0.11** | **0.85** | **1.27** |
| CFU_*L.m.* : Temp.25 | -0.11 | 0.14 | -0.37 | 0.16 |
| CFU_*L.m.* : Temp.30 | -0.10 | 0.26 | -0.60 | 0.41 |
| **CFU_*L.m.* : Temp.HW** | **0.39** | **0.16** | **0.07** | **0.70** |
| Random effect (batch) | 0.74 | 0.41 | 0.31 | 1.83 |
